# Quantification of circadian interactions and protein abundance defines a mechanism for operational stability of the circadian clock

**DOI:** 10.1101/2021.08.27.456017

**Authors:** James S. Bagnall, Alex A. Koch, Nicola J. Smyllie, Nicola Begley, Antony Adamson, Jennifer L. Fribourgh, David G. Spiller, Qing-Jun Meng, Carrie L. Partch, Korbinian Strimmer, Thomas A. House, Michael H. Hastings, Andrew S. I. Loudon

## Abstract

The mammalian circadian clock exerts substantial control of daily gene expression through cycles of DNA binding. Understanding of mechanisms driving the circadian clock is hampered by lack of quantitative data, without which predictive mathematical models cannot be developed. Here we develop a quantitative understanding of how a finite pool of BMAL1 protein can regulate thousands of target sites over daily time scales. We have used fluorescent correlation spectroscopy (FCS) to track dynamic changes in CRISPR-modified fluorophore-tagged proteins in time and space in single cells across SCN and peripheral tissues. We determine the contribution of multiple rhythmic processes in coordinating BMAL1 DNA binding, including the roles of cycling molecular abundance, binding affinities and two repressive modes of action. We find that nuclear BMAL1 protein numbers determine corresponding nuclear CLOCK concentrations through heterodimerization and define a DNA residence time of 2.6 seconds for this complex. Repression of CLOCK:BMAL1 is in part achieved through rhythmic changes to BMAL1:CRY1 affinity as well as a high affinity interaction between PER2:CRY1 which mediates CLOCK:BMAL1 displacement from DNA. Finally, stochastic modelling of these data reveals a dual role for PER:CRY complexes in which increasing concentrations of PER2:CRY1 promotes removal of BMAL1:CLOCK from genes consequently enhancing ability to move to new target sites.

## Introduction

The 24-hour light-dark cycles inherent to our planet have led to the evolution of molecular circuits capable of conveying time of day information, commonly known as circadian clocks. In mammals, cell-autonomous circadian clocks operate in virtually all cells across tissues and enables coordination of numerous biological processes, including metabolism, immunity, and cell cycle progression (1, 2).

Autonomous cellular clocks are characterized by transcription-translation feedback loops (TTFLs), leading to cycles in protein and mRNA tuned to the 24-hour rhythms of the day-night cycle. Central to the mammalian circadian clock is the heterodimeric transcription factor comprised of CLOCK (circadian locomotor output cycles protein kaput) and BMAL1 (brain and muscle ARNT-like 1) that searches the genome to bind consensus sequence E-box sites (CANNTG), inducing expression of several hundred clock-controlled output genes every day. Targets include key circadian negative feedback regulators, *Period* (*Per1*, *Per2*, *Per3*), *Cryptochrome* (*Cry1* and *Cry2*) and a secondary loop regulated by *Nr1d1* and *Nr1d2* (3–6). These proteins act to repress the activity of CLOCK:BMAL1 to form a delayed negative feedback loop driving daily oscillations. In a current model, it is proposed that PER and CRY proteins dimerize to form a repressive complex with CK1 (casein kinase 1) to promote the removal of CLOCK:BMAL1 from DNA and thereby repress transactivation of target genes, while CRY1 independently binds the PAS domain core of CLOCK:BMAL1 and the BMAL1 transactivation domain leaving DNA binding intact whilst repressing recruitment of additional transcriptional coactivators (7, 8). An additional feedback loop is conferred by the protein REV-ERBα, which operates as a transcriptional repressor of *Bmal1*, resulting in a cycle of BMAL1 protein abundance (6).

Ultimately, a prerequisite for generation and output of cellular circadian rhythms is the ability of a finite pool of CLOCK:BMAL1 heterodimer protein to bind rhythmically to specific target sequences leading to the regulation of circadian gene expression in a specific cell type (9). Currently, we have very little insight into the quantitative biology of this process. Heterodimeric formation of transactivating and repressive complexes is a well-defined feature of the circadian molecular circuit, including the formation of CLOCK:BMAL1 and PER:CRY complexes (3, 7, 8). Recently, PER:CRY proteins have been described as part of very large macromolecular complexes within the cell (10). We have previously visualized several core circadian proteins, and from this measured the spatiotemporal profile and protein abundance for BMAL1 and PER2 (11, 12). PER2 was found to cycle with a maximum amplitude of 12,000 copies per cell in fibroblasts and without circadian gating of nuclear localization, contrary to observations in the *Drosophila* clock (11, 13). A recent study using a cancer cell line model has also shown that CRY1 protein remains nuclear at all circadian phases and at markedly higher abundance than its partner protein PER2 (14).

In order to gain insight into the operation of core circadian clock proteins, we generated a genetically modified mouse in which CRY1 has been C-terminally fused with a fluorescent protein. We crossed this line to a previously described strain of mice expressing fluorescent-tagged BMAL1. We then used advanced imaging in both ectopically transformed cell lines and endogenously modified mice to characterize governing parameters in the regulation of BMAL1 DNA binding, including repression by PER2 and CRY1. We constructed mathematical models of the complex interactions and phased timings from multiple molecular species and experimentally inaccessible complexes, demonstrating how DNA binding in the peripheral circadian clock is regulated.

Using a combination of mathematical modelling and experimental validation, our data reveal that high affinity interactions between circadian protein complexes serve to offset the low abundances of circadian proteins. In this way, the abundance of key components of the molecular clockwork is positioned optimally to regulate E-box binding. This is partly facilitated through PER2:CRY1 mediated displacement of CLOCK:BMAL1, such that PER2 protein serves a dual role, acting as both a component of the negative feedback arm but also to redistribute CLOCK:BMAL1 to new target sites. Thus, the stochiometric balance of PER:CRY with CLOCK:BMAL1 is critical for the elucidation of the full cellular circadian repertoire.

## Results

### BMAL1 determines nuclear localization and mobility of CLOCK

To quantify the properties of BMAL1 and CLOCK proteins, we first used NIH/3T3 fibroblasts expressing fluorescent fusion proteins via a ubiquitin ligase C promoter, delivered by lentiviral transduction either singly (LV1) or as two sequential transductions (LV2) (**Fig. 1A**) (15). Expression of the transgene was in >10-fold excess over the native unfused protein, as determined by single molecule Fluorescence In Situ Hybridisation (**Appendix SI, Fig. S1**). Confocal microscopy of tagRFP::CLOCK or BMAL1::EGFP showed BMAL1 expression to be strongly localized to the nucleus whereas CLOCK was predominantly cytoplasmic when expressed alone (**Fig. 1A**). Co-expression of both proteins in the same cells caused localization of tagRFP::CLOCK to the nucleus, in agreement with earlier studies which showed cytoplasmic CLOCK localization in BMAL1-deficient cells, and that circadian regulation of nuclear localization of CLOCK correlated with BMAL1 availability (16, 17). We also transduced cells with a fluorescent fusion of a DNA binding mutant of BMAL1, in which a leucine is substituted for glutamic acid in the basic helix-loop-helix region of the protein; referred to as L95E. This protein re-localized tagRFP::CLOCK protein to the nucleus from the cytoplasm in an manner equivalent to WT BMAL1 (**Fig. 1A**) (3).

**Figure 1.**
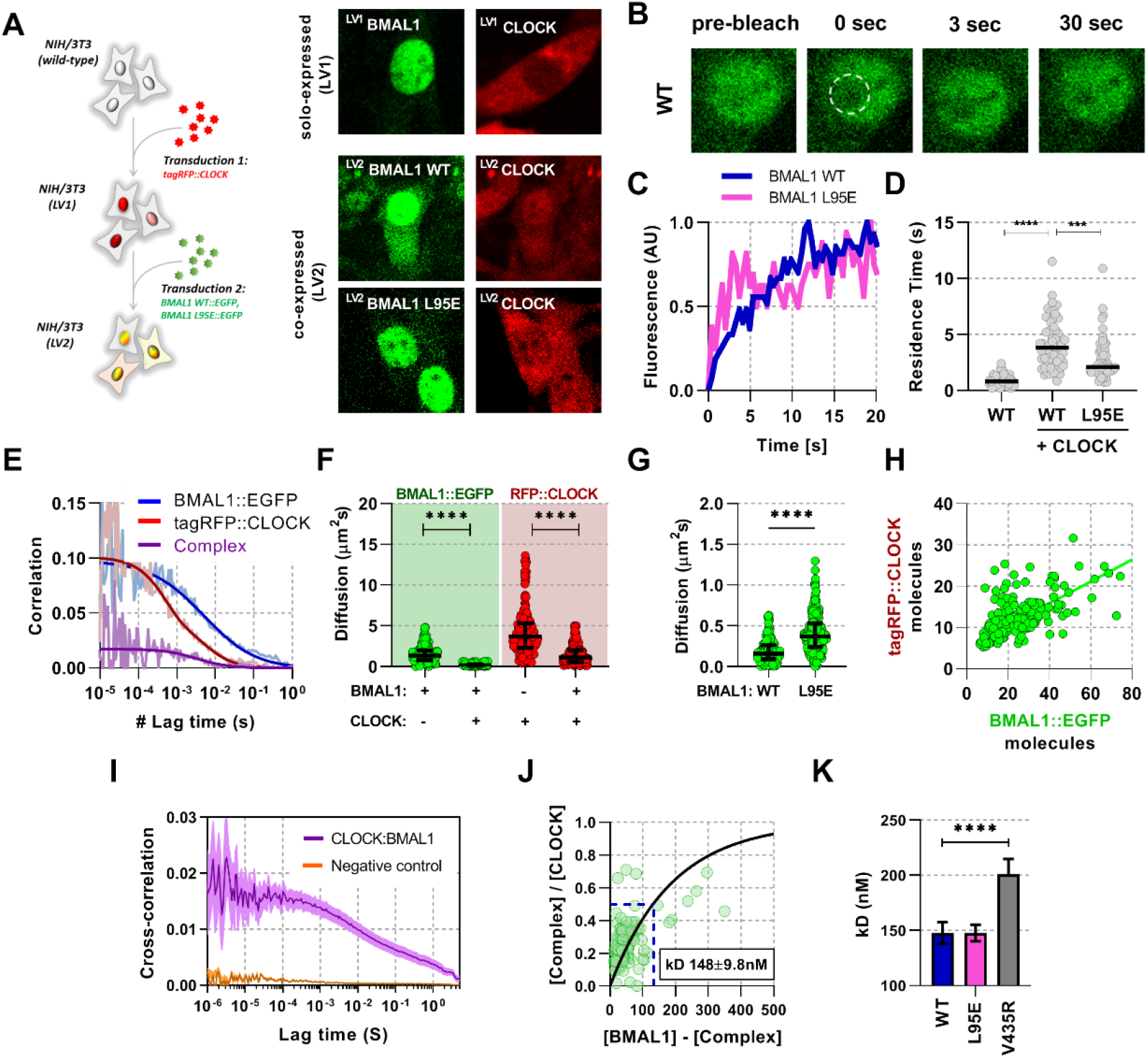
BMAL1 and CLOCK mobility is regulated by dimerization and DNA binding. (**A**) NIH/3T3 cells are either singularly or sequentially transduced to express fluorescent fusions with CLOCK or BMAL1 (wildtype and mutant variants). Confocal microscopy images of cells solo-expressing (LV1) either tagRFP::CLOCK or BMAL1::EGFP or co-expressing (LV2) them together (including BMAL1L95E DNA binding mutant). (**B**) Confocal microscopy images for photobleaching of ^LV2^BMAL1::EGFP-RFP::CLOCK labelled cells, either with wild-type or BMAL1L95E DNA binding mutant. Images show nuclei and highlight region of bleaching. (**C**) Representative fluorescence recovery curves of bleach region for B. following normalisation. (**D**) Residence time calculated as the inverse of k_OFF_ (s^-1^), determined from fitting the recovery data in C. with a single component binding model (n = 69, 58 and 51 cells). Bar represents median values. (**E**) Representative auto- and cross-correlation data showing raw data and fit lines for monomeric and complexed fluorescent proteins. (**F**) FCS data showing diffusion for BMAL1 and CLOCK in solo- and co-expressed conditions (n = 173, 152, 198 and 185 cells). (**G**) FCS results for BMAL1::EGFP diffusion for NIH/3T3 cells that co-express tagRFP::CLOCK. Data shown is for comparison of BMAL1 as either wild-type of L95E DNA-binding mutant. Bars show median and interquartile range. (**H**) Correlation of nuclear protein quantification showing relationship of BMAL1::EGFP with tagRFP::CLOCK for both wildtype and DNA binding mutant (n=221 cells from 3 biological replicates). (**I**) Average cross-correlation curves for BMAL1::EGFP (WT) with tagRFP::CLOCK (n = 140) compared to a non-interacting control of NLS::EGFP co-expressed with tagRFP::CLOCK (n = 408). (**J**) Dissociation plot from FCCS data for BMAL1::WT and tagRFP::CLOCK. (**K**) Summary of calculated dissociation constants across all conditions, including BMAL1 dimerization mutant, V435R (n= 156, 274 and 244). Mann-Whitney non-parametric test to determine significance (values are denoted as p>0.05 ns, p<0.05 *, p<0.01 **, p<0.001 *** and p<0.0001 ****)

We next performed Fluorescence Recovery After Photobleaching (FRAP) experiments to test the impact of CLOCK on the recovery dynamics of a bleached nuclear region of BMAL1::EGFP, by comparing responses with or without co-expressed tagRFP::CLOCK (**Fig. 1B-C**). BMAL1 recovery half-life was found to be insensitive to the diameter of the bleached region, indicating that binding contributes to the recovery profile rather than this being a solely diffusion-led process (**Appendix SI, Fig. S2**) (18). Reaction binding equations were fitted to determine the rate of dissociation, K_OFF_, for BMAL1::EGFP, the reciprocal of which equates to an average characteristic duration of binding or residence time. Residence time of BMAL1::EGFP was increased in the presence of tagRFP::CLOCK, consistent with a requirement for CLOCK to bind DNA (**Fig. 1D**). The mean residence time for the fluorescent CLOCK:BMAL1 complex was 4.13 seconds (95% CI, 0.57), a value consistent with DNA residence times for similar transcription factors (16, 19). Using the L95E DNA-binding mutant protein, we saw significantly reduced residence time of 2.83 seconds (95% CI, 0.54). Notably, the initial publication of the BMAL1 L95E mutant showed a 2-fold reduction in PER2::LUC expression and so suggests a strong relationship between DNA binding and transcriptional output (3).

To investigate this further we used Fluorescence (Cross) Correlation Spectroscopy (F(C)CS) (20), which enabled determination of live cell concentration and diffusion properties of individually fluorescent-labelled BMAL1 and CLOCK proteins, and their interactions when co-expressed (**Fig. 1E).** A normal diffusion model fitted the majority of data collected from cells expressing free EGFP or NLS::EGFP proteins, as previously reported (21), whereas anomalous diffusion models – sub-diffusion caused by a range of factors such as DNA interactions and molecular crowding – accounted for a >20% fraction, which in this instance may be explained by molecular crowding (22). In comparison, for the fusion proteins, anomalous diffusion models accounted for >80% of all BMAL1 data sets (**Appendix SI, Fig. S3**). We used an anomalous diffusion model for all further analyses of circadian proteins to calculate diffusion coefficients and protein concentrations.

Singly expressed fluorescent BMAL1 and CLOCK were found to diffuse rapidly with a median coefficient of 9.2 μm^2^/s (SD, 3.3) and 12.6 μm^2^/s (SD, 6.1), respectively. In contrast, co-expression significantly reduced the rate of diffusion to 1.9 μm^2^/s (SD, 1.3) and 4.7μm^2^/s (SD, 3.2) for BMAL1 and CLOCK respectively (**Fig. 1F**). The L95E mutant diffused more rapidly than WT BMAL1, consistent with fewer interactions in the nucleus (**Fig. 1G**). When co-expressed, WT BMAL1 and CLOCK exhibited a 2:1 concentration ratio in the nucleus (**Fig. 1H, Appendix SI, Fig. S3**). F(C)CS was then used to observe this interaction and determine a live-cell dissociation constant (K_D_; reciprocal measure of affinity) (23). A positive cross correlation curve was observed between BMAL1::EGFP and tagRFP::CLOCK that was not apparent in cells expressing NLS::EGFP with tagRFP::CLOCK (**Fig. 1I**). To calculate K_D_, we fitted a one-site saturating binding curve to the relationship between heterodimer and monomer (**see supplementary material**) which yielded a value of 148 nM (SD, 9.8) for WT BMAL1::EGFP and tagRFP::CLOCK (**Fig. 1J**). The K_D_ was measured for cells with the reverse fluorescent protein labelling, namely EGFP::CLOCK and BMAL1::tagRFP, finding similar a value of 145 nM (SD, 4.8), although a stronger interaction was found in vitro by surface plasmon resonance (**Appendix SI, Fig. S4**). Previous work found that the V435R mutation of BMAL1 in the PAS-B domain, leads to reduced dimerization with CLOCK (3). We used the V435R mutation to confirm our F(C)CS measurements by co-expressing V435R-BMAL1 and WT-CLOCK. This elicited a ~1.5-fold reduction in interaction affinity, resulting in a K_D_ of 201 nM (SD, 14) (**Fig. 1K**) and a reduction from 2:1 to a 4:1 ratio of BMAL1 and CLOCK in the nucleus (**Appendix SI, Fig. S4E**). In contrast, the BMAL1 L95E DNA binding mutant showed no difference in interaction affinity compared to WT BMAL1 protein. These data demonstrate that BMAL1 is a critical determinant of the localization, mobility and concentration of CLOCK in the nucleus.

From this, we can infer the abundance of nuclear CLOCK from measurements of BMAL1, and make use of available endogenously labelled Venus::BMAL1 mice to measure remaining DNA binding parameters. First, we confirmed our existing cell line measurements of binding rates and diffusion using the Venus::BMAL1 mice (12), and showed that they remain within a similar range across a number of primary cell types, including macrophages and pulmonary fibroblasts (**Appendix SI, Fig. S5**). We also measured protein number of the endogenous BMAL1, finding that total copies per nucleus vary from 1000-10000 between individual cells, likely due to desynchrony and differing nuclear volumes. Moreover, a large overlap in nuclear copy numbers was observed across all cell types despite substantial changes in the mean. These data are critical in our understanding of the ratio of BMAL1 to target sites, effectively determining the capacity to regulate the full repertoire of target genes within a specific cell type.

### Quantification of strong rhythmic interaction of BMAL1 with CRY1

The ability to measure BMAL1 amounts to infer copy number of CLOCK, allows us to measure other critical interactions with BMAL1. This includes the repressive action of CRY1 binding to CLOCK:BMAL1, resulting in reduced transactivation. To explore the interaction between BMAL1 and CRY1, we generated a genetically modified mouse in which CRY1 has been C-terminally fused with the red fluorescent protein mRuby3 using CRISPR-mediated genomic editing to insert the coding sequence, replacing the endogenous CRY1 stop codon (**Fig. 2A**) (24, 25). First, to test any potential impact on circadian pace-making, we measured CRY1::mRuby3 fluorescence in whole-field organotypic SCN slices (**Fig. 2B**) which exhibited regular cycles in red fluorescence with a period of 23.9 hours (SD, 0.6) (**Fig. 2C**) (11). Additionally, wheel running measurements of these mice confirmed normal behavioral rhythmicity (**SI appendix Fig. S6**). We next crossed these mice to the Venus::BMAL1 mouse line (12), previously inter-crossed with a PER2::LUCIFERASE background (26) (**SI appendix Fig. S5**) to provide an independent circadian phase-reference marker (referred to as BMAL1xCRY1 labelled mouse). Using isolated lung fibroblasts from BMAL1 x CRY1 mice we assessed bioluminescence in response to dexamethasone (DEX) synchronization, and observed 23.3 hour cycles (SD, 0.6) which were sustained for >4 days (**Fig. 2D-E**). From this, we are confident that the fluorescent fusion proteins do not disrupt the normal operation of the circadian pacemaker.

**Figure 2.**
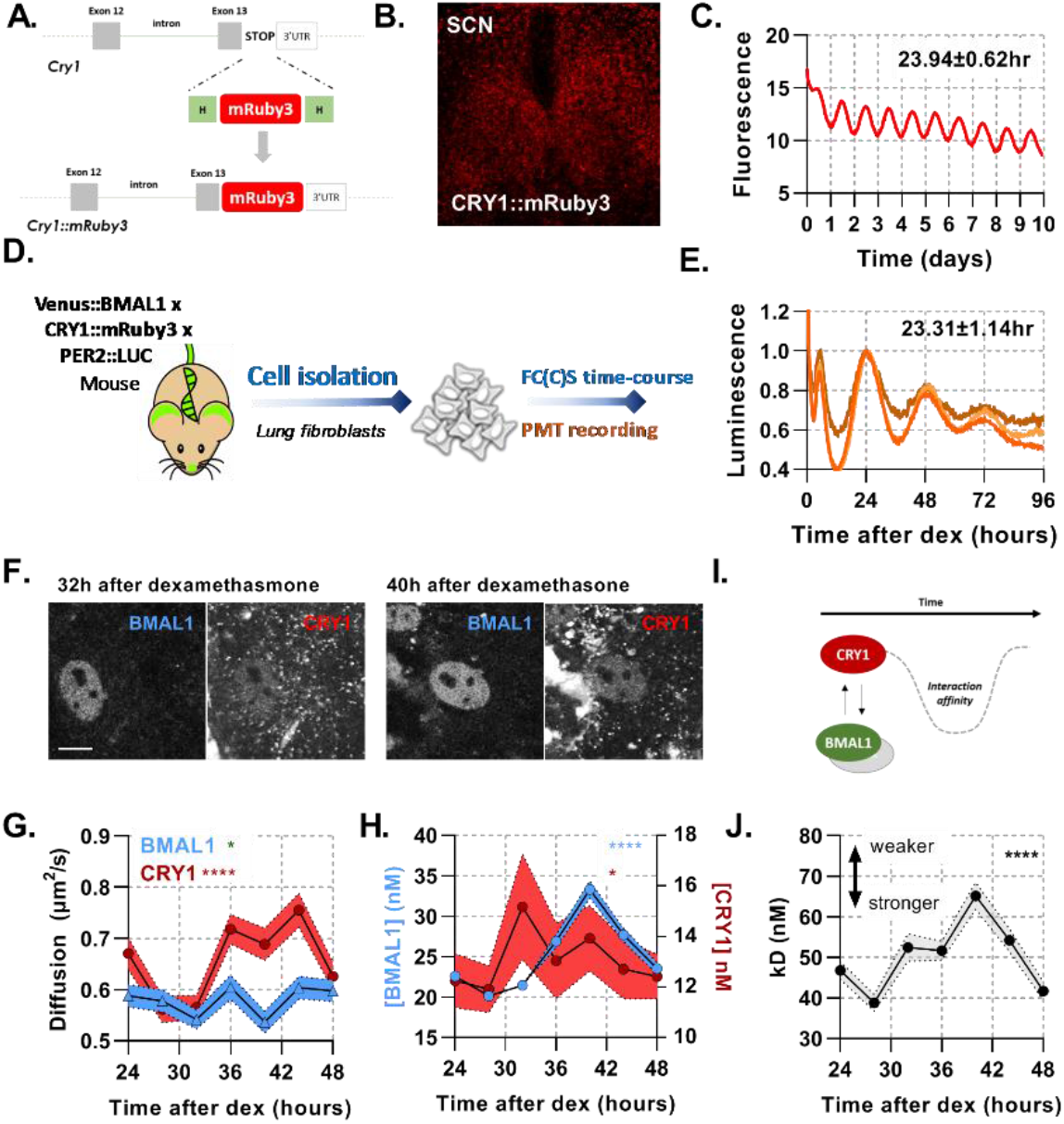
A rhythmic and strong interaction observed between BMAL1 and CRY1 facilitates repression. (A) Schematic representation of newly made transgenic mouse engineered to express CRY1::mRuby3, Venus::BMAL1 and PER2::LUC. (B) Confocal microscopy image of SCN organotypic slice expressing CRY1::mRuby3. (C) Quantification of mRuby3 fluorescence over time for the whole SCN, with mean and standard deviation of period. (D) Experimental set up to measure isolated primary lung fibroblasts from Venus::BMAL1 x CRY1::mRuby3 x PER2::LUC labelled mice synchronised with dexamethasone. Parallel cell cultures were analysed for luminescence and also by FCS over a time-course, measured every 4 hours. (E) Luminescence recordings of isolated primary lung fibroblasts from BMAL1 x CRY1 x PER2 labelled mice synchronised with dexamethasone. Data shown is for three independent replicates, with mean and standard deviation of period shown. (F) Confocal images of two cells shown for Venus::BMAL1 and CRY1::mRuby3 over time. FCS determined measurement for diffusion coefficient (G) and protein concentration (H) of Venus::BMAL1 and CRY1::mRuby3 (n = 136, 143, 173, 131, 158, 121 and 132; line shows the mean and error envelopes show the SEM). (I-J) Interaction strength between BMAL1 and CRY1 was also measured over time as illustrated by the schematic of affinity as well as plotted values of dissociation constant (error envelope shows the standard deviation).

Using the same synchronization approach, we then measured BMAL1xCRY1 fluorescence in single cells every 4 hours from 24 to 48 hours post-DEX synchronization, using F(C)CS (**Fig. 2F**). Both fluorescent signals were localized to the nucleus. Venus::BMAL1 showed a consistent diffusion pattern over a circadian cycle, with a mean diffusion coefficient of 0.58 μm^2^/sec (SD, 0.03), whereas CRY1 mobility exhibited circadian variance, with slow diffusion 28 hours post-DEX and elevated diffusion rates 12 hours later (**Fig. 2G**). Interestingly, this change in mobility is consistent with a binding to a mass equivalent to the molecular weight of PERIOD2 (**see supplementary material**). Peak protein concentrations of BMAL1 and CRY1 had an approximate and appropriate phase-separation of 8 hours (27). Auto-correlation analyses revealed the concentration of BMAL1 is on average 1.92 fold (SD, 0.32) higher than CRY1, with a mean concentration of 29.3 nM and 13.4 nM respectively (equating to approximately 16,000 and 7,000 molecules per nucleus), consistent with the range we reported earlier for PER2 (11). The amplitude of CRY1 was found to be shallow, cycling from 7.9 molecules per confocal volume (cv, FCS measurement volume) at T28 to 10.0 molecules per cv at T32, comparable to the approx. 25% amplitude observed for CRY1 in the SCN (**Fig. 2C**). BMAL1 demonstrated a larger amplitude, cycling from 9.8 molecules per cv at T28 to a peak of 20.2 molecules per cv (**Fig. 2H**).

We then analyzed the interaction affinity between BMAL1 and CRY1 over time (**Fig. 2I**). This interaction exhibited significant changes over a 24-hour cycle, with the strongest interaction at T28, K_D_ = 38.8 nM (SD, 2.1), and weakest at T40, K_D_ = 65.1 nM (SD, 3.4) (**Fig. 2J**). These profiles were found to correlate with the diffusion profile of CRY1 (**Fig. 2G**). Interestingly, the mean interaction strength between BMAL1 and CRY1 is >2-fold stronger than that between BMAL1 and CLOCK (**Fig. 1K**). A similar relationship was found *in vitro* when measuring interactions using biolayer interferometry (28). This is therefore consistent with a model in which the low abundance of the CRY1 repressor is offset by a high affinity for the CLOCK:BMAL1 heterodimer.

The changes in the diffusion profile of CRY1 are consistent with its association with an additional binding partner, such as PER2, thereby altering the affinity of CRY1 for CLOCK:BMAL1 (28, 29). To measure the interaction between CRY1 and PER2 directly, we transduced NIH/3T3 cells with lentivirus so that cells constitutively express EGFP::PER2 or CRY1::tagRFP fusion proteins. In both cases, the expressed protein was found to localize predominately to the nucleus, although some cytoplasmic fluorescence was observed. When co-expressed, subcellular localization was unchanged, although large punctate aggregates of signal were observed (**Fig. 3A**). PER2 was found to have the slowest diffusion coefficient recorded within all our F(C)CS measurements, when in the non-aggregate space. PER2 mobility was not altered following co-expression with CRY1, whereas CRY1 exhibited reduced mobility in the presence of ectopic EGFP::PER2 (**Fig. 3B**). The diffusion coefficient for CRY1 co-expressed with PER2 was similar to measurements of the endogenous protein (**Fig. 2G**), suggesting PER2 and CRY1 exhibit similar stoichiometry within these cells. The anomalous diffusion model fit the majority of data sets, including ^LV1^CRY1, ^LV1^PER2 and ^LV2^PER2. However, normal diffusion models accounted for >50% of ^LV2^CRY1 correlation analyses suggesting a distinct change in CRY1 following interaction with PER2, potentially from a loss of significant DNA binding of the CLOCK:BMAL1 complex (**Appendix SI, Fig. S7**). Best fit models for each data set demonstrated a strong affinity between PER2 and CRY1 with a K_D_ of 81.8 nM (SD, 4.9) (**Fig. 3C**) and consistent with previous in vitro measurements (30).

**Figure 3.**
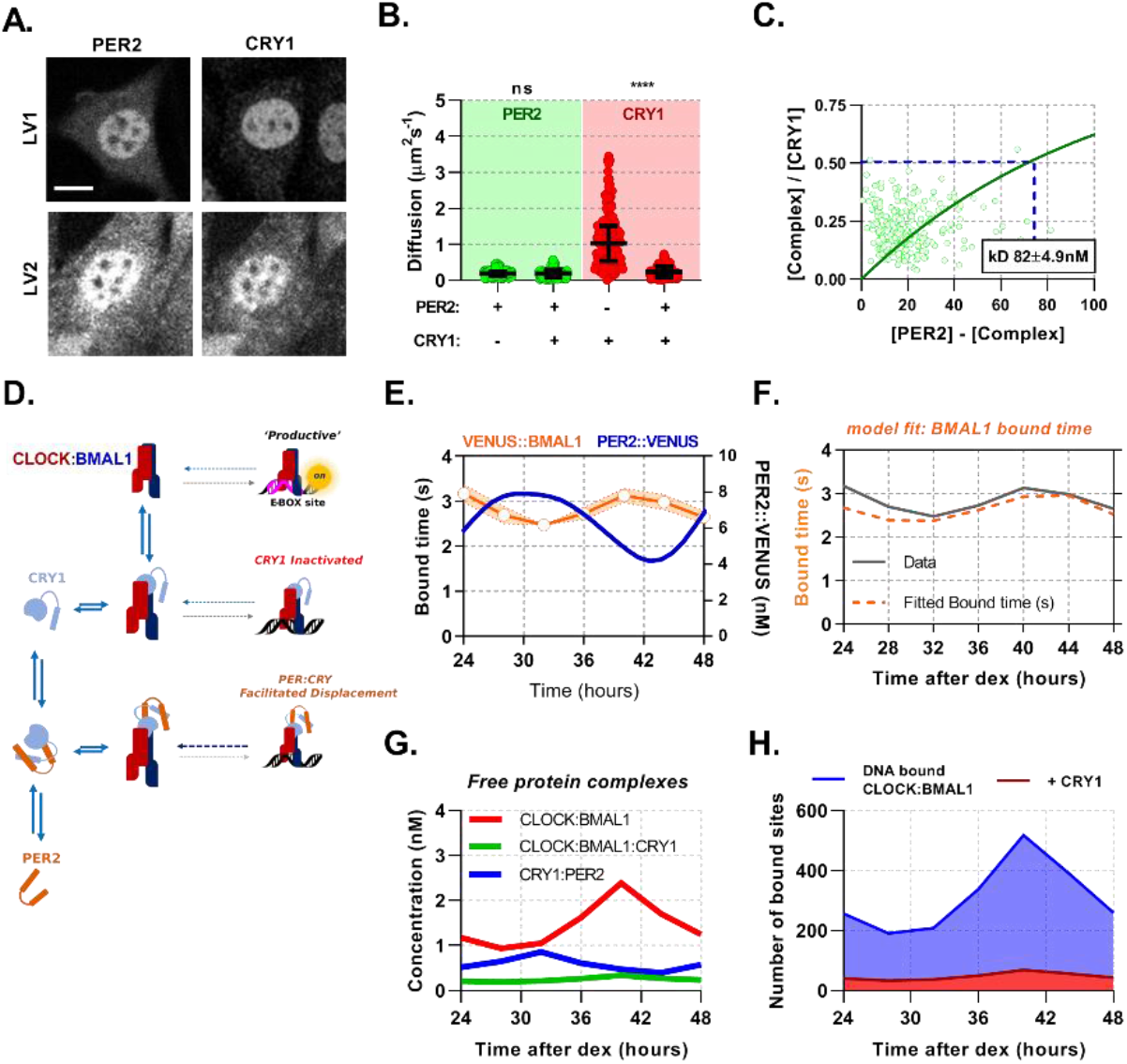
PER2 acts via CRY1 to mediate rhythmic displacement of CLOCK:BMAL1 from DNA. **(A)** Confocal images of transduced NIH/3T3 cells that either solo- or co-express PER2 and CRY1. (B) FCS data showing diffusion for PER2 and CRY1 in solo- and co-expressed conditions (n = 165, 174, 274 and 274 cells). (**C**) Dissociation plot from nuclear FCS measurements for EGFP::PER2 and CRY1::tagRFP (n=274). **(D)** Schematic representation of model topology used for the deterministic model of CLOCK:BMAL1 DNA binding. **(E)** Primary lung fibroblasts from BMAL1 x CRY1 x PER2 mice were synchronised with dexamethasone. Plot shows PER2 concentration as measured via FCS by Smyllie et al 2016 (11) as well as mean BMAL1 binding time (showing SEM error envelope). Binding time was measured by confocal FRAP measurements performed on the Venus::BMAL1 fluorescence. Orange line shows the inverse of k_OFF_ (s^-1^), determined from fitting the recovery data with a single component model (n = 48, 70, 82, 63, 82, 64 and 65 cells). **(F)** ODE model was fit to FRAP binding data from E. and using a measured input for PER2 nuclear concentration previously determined in Smyllie. et al 2016. Model output showing (**G**) inferred nuclear concentrations for molecular complexes (**H**) and CLOCK:BMAL1 without and with CRY1 bound to target sites (see supplementary materials for parameters).

Quantitative data are an enabling and often essential component of stringent mathematical modelling of cell signaling (31). Having quantified the necessary parameters, we then sought to use them in developing a mathematical model of CLOCK:BMAL1 DNA binding, with the aim of understanding how the multiple regulatory motifs of changing molecular concentrations, interactions and binding kinetics coalesce to regulate DNA binding and transcriptional activation of BMAL1. We modelled the system using a set of ordinary differential equations (ODEs) to depict a current understanding of the system; BMAL1 dimerizes with CLOCK, which may subsequently bind and unbind to DNA target sites. To model repression, CRY1 may either inactivate CLOCK:BMAL1 via direct binding or, via dimerization to PER2, form PER2:CRY1 (mimicking complexes with CK1) to displace CLOCK:BMAL1 from DNA (**Fig. 3D**) (3, 7–9). The latter would presumably lead to rhythmic changes in the residence time of BMAL1 and provide a sensible option to fit and complete the model.

To assess dynamic changes in inferred DNA binding rates of BMAL1, we isolated lung fibroblasts from BMAL1xCRY1 mice and determined the k_OFF_ values by FRAP following DEX synchronization. Measurements of BMAL1 protein recovery were taken every 4 hours from 24 h to 48 h post-DEX, showing k_OFF_ to be rhythmically regulated (**Fig. 3E**). The BMAL1 k_OFF_ profile was in antiphase to recordings of nuclear PER2 concentrations from Smyllie et al. 2016 (**Fig. 3E**) (11). To fit all parameters to the model (supplementary material and Table S1), the measured concentrations of PER2 (11), CRY1 and BMAL1 were used as inputs, using data described in Figures 2H and 3F. On/Off rates were constrained to measured dissociation constants from F(C)CS (**Appendix SI, Table S1**), with the K_D_ value between BMAL1 and E-box sites set at 10 nM, as measured by Huang *et al* (3). Using a mean value from multiple published ChIP-Seq data, the potential number of DNA target sites was set as 3436 (**Appendix SI, Table S1**) (9, 32).

The ODE model was then fitted by simulating FRAP so that a model-derived k_OFF_ could be used against our experimental data (**Fig. 3E**) via Chi^2^ minimization (Chi^2^, 7.53) (**Fig. 3F, SI Table S2**). Next, we used this model to infer experimentally inaccessible complexes, specifically PER2:CRY1, CLOCK:BMAL1 and CLOCK:BMAL1:CRY1 (**Fig. 3G**). We find that free CLOCK:BMAL1 (unbound to DNA) cycles in remarkably low abundance from 0.9 nM to 2.4 nM, which equates to a change of ca. 809 molecules, similar to that of the PER2:CRY1 complex. Furthermore, predicted DNA binding of CLOCK:BMAL1 has an average baseline of 190 sites bound at any one time, rising to 518 sites at the peak, in agreement with the expected ~2-fold peak enrichment from ChIP reports (9, 32) (**Fig. 3H**). The model also suggests ~50% of the available transcription factor complex is engaged with site-specific interactions with availability predominantly limited by the K_D_ with CLOCK. Finally, DNA bound CLOCK:BMAL1:CRY1 complex was persistent, with low abundance cycling from 32 to 40 target sites (accounting for 14% of total site bound BMAL1).

### Circadian proteins are within an optimal expression range to modulate E-Box binding

The topology of the circadian molecular circuit is preserved across all cell types, yet it is also known that different cell types have widely differing repertoires of target genes and accessible genomic target sites for CLOCK:BMAL1 to bind (32). We therefore pursued the extent to which varying the number of target sites may have an impact on the available pool of CLOCK:BMAL1 to bind target sequences, as calculated by site occupancy (the % sites occupied at any given moment). We simulated the model over a biological range of binding sites (1,000 – 10,000), informed by multiple BMAL1 and CLOCK ChIP data sets (**Fig. 4A**) (9, 32). We found target site occupancy decreased marginally from 16.2% to 13.6%, showing that any variance between different numbers of available target sites has minimal impact. We then explored how varying binding parameters affected site occupancy, relating them to the variability observed in our data sets but considering values beyond these limits. The CLOCK:BMAL1 unbinding rate accounted for a 21.5% change when residence times across the observed physiological range are considered (**Fig. 4B**); outside of this range occupancy begins to saturate so that a further 30 second increase in residence time only accounts for an additional 12.5% binding. Therefore, the unbinding rate, as measured experimentally, is optimally positioned to regulate target site occupancy in a manner consistent with the displacement mechanism governed by PER:CRY.

**Figure 4.**
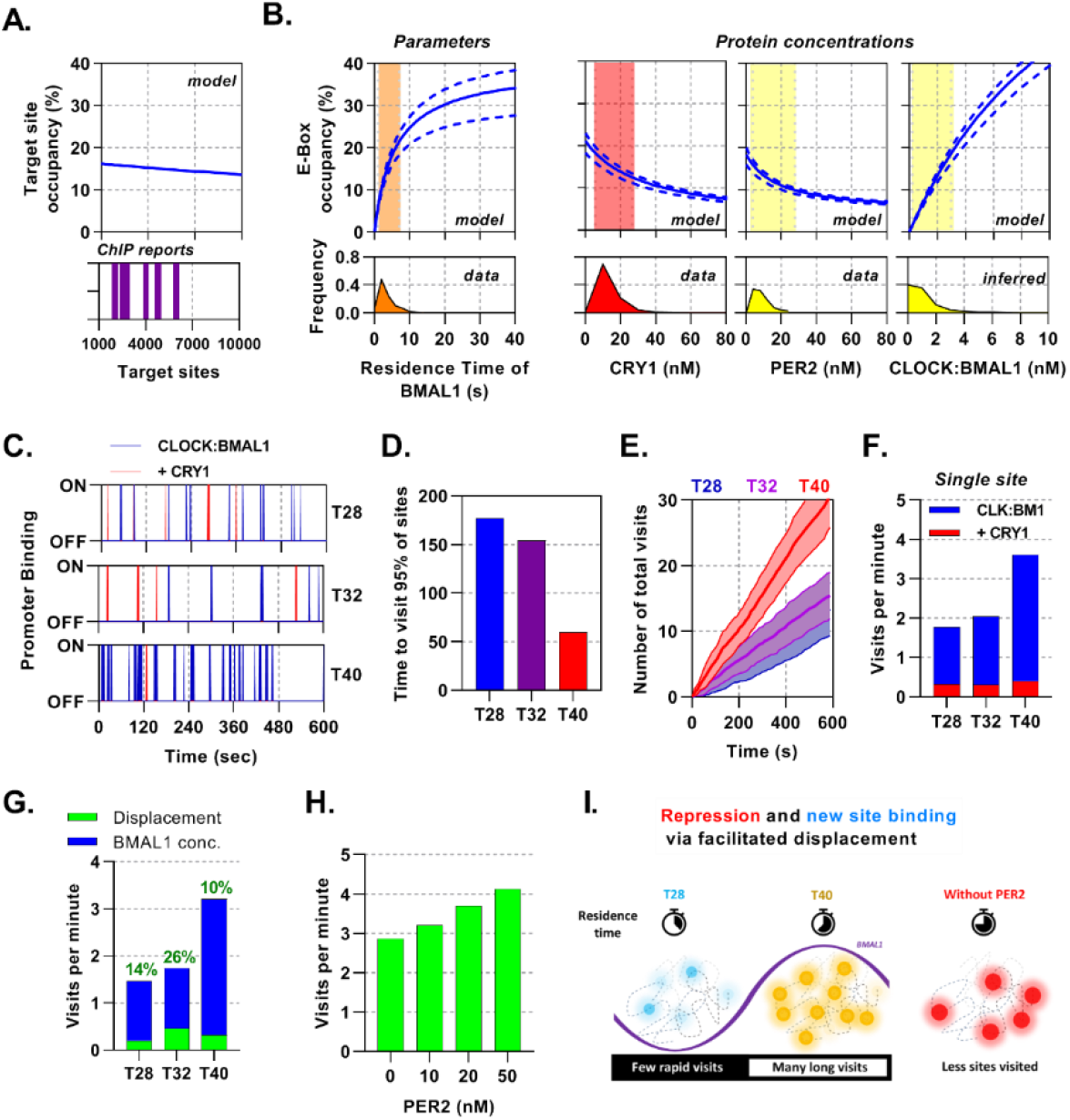
Mathematical modelling demonstrates dual function of PER:CRY mediated repression. **(A-B)** Sensitivity analysis of the deterministic binding model showing relationship of measured parameters (bottom) against model for occupancy of active BMAL1:CLOCK on target sites (top). **(A)** Changing number of target sites with data matched to BMAL1 ChIP data sets. **(B)** From left to right, the effect of changing residence time of CLOCK:BMAL1, or protein concentrations. Histograms show measured concentrations for corresponding proteins across all conditions/cells. The 10^th^ to 90^th^ percentile is highlighted. **(C-G)** Stochastic binding model outputs using parameters corresponding to T28, T32 or T40 post dexamethasone BMAL1 x CRY1 data sets. **(C)** Shows a promoter corresponding to the average binding rate of CLOCK:BMAL1, **(D)** the time to visit 95% of target sites once and (E) number of visits to a single promoter over time. **(F)** Average number of visits per minute to a target site showing active and CRY1 repressed CLOCK:BMAL1 visits. **(G)**. Comparison of the contribution of BMAL1 concentration (blue) and PER2 facilitated displacement (green) on the visits per minute to a target site. Percentage contribution indicated. **(H)** Relationship of PER2 protein concentration to site visitations per minute by CLOCK:BMAL1 using parameters for T40 explored over different concentrations of PER2. **(I)** The action of CRY:PER leads to short-lived transient binding of CLOCK:BMAL1 to DNA, working as both a repressive action whilst also facilitating binding to new target sites.

Additionally, protein concentrations vary across circadian time as well as individual cells and cell types (**Appendix SI, Fig. S5)**. We therefore simulated target-site occupancy across varied biologically plausible concentrations for CRY1, PER2 and CLOCK:BMAL1 and calculated the fraction of occupied sites (**Fig. 4B**). Increasing CRY1 and PER2 led to a reduction in target-site occupancy, whereas a rise in CLOCK:BMAL1 led to a substantial increase and in both cases. Moreover, the biologically observed range occupied the most sensitive part of the curve, such that oscillations in protein copy number can evoke significant changes of occupancy. Hence, the system is positioned to make efficient use of the biological concentrations of the constituent proteins.

### Mathematical modelling demonstrates dual function of PER2:CRY1 mediated repression

Site occupancy is a function of the average residence time of transcription factors bound to DNA; consequently, highly frequent and short events would appear the same as infrequent and long binding events. To infer these masked kinetics, which are obscured in our mean based ODE model, we use a stochastic binding model to simulate individual molecules of CLOCK:BMAL1 binding to target DNA sites in a well-mixed system (33). For our simulations, we have used the average number of molecules and effective dissociation rates determined for T28, T32 and T40 hours post-DEX for lung fibroblasts (**Fig. 4C**) arising from our previously described ODE model. T28 and T40 represent trough and peak of BMAL1 (**Fig. 2H**) respectively, whereas T32 and T40 represent the trough and peak of PER:CRY protein amounts (**Fig. 3G**).

Alongside binding of active CLOCK:BMAL1, we also considered target sites bound by CLOCK:BMAL1:CRY1, which are thought to be transcriptionally inactive whilst also blocking target site access to active molecules. At T40, when there is the maximum amount of CLOCK:BMAL1, 95% of the 3436 target sites would be bound at least once within a minute, changing to 3 minutes at T28 (**Fig. 4D**), contributing towards a small degree of heterogeneity. From the perspective of a single promoter at T40 there are ~3.1 visits per minute by CLOCK:BMAL1, which is reduced down to ~1.3 visits per minute at T28 (**Fig. 4E**), with further reductions in individual cells with lower concentrations of CLOCK:BMAL1 protein (**Appendix SI, Fig. S8)**. We then separated the total visits per minute into those occurring as CLOCK:BMAL1 compared to those occurring as the CLOCK:BMAL1:CRY1 complex, finding the latter to remain relatively persistent across time points and making up ~15% of total visits, mirroring results for our ODE model. Our stochastic model therefore predicts that oscillating amounts of BMAL1 and CRY1 protein amounts, as well as the changing interaction affinity, may actually help preserve the concentration of CLOCK:BMAL1:CRY1 target binding events across circadian time (**Fig. 4F**).

Repression of CLOCK:BMAL1 activity by CRY1 requires continuous interaction and hence is limited by concentration. We hypothesized that this would be different for the PER2:CRY1-mediated displacement of CLOCK:BMAL1 from target DNA sites. To test this, we first investigated how the number of visits per minute would be affected by clamping the input values of K_OFF_ and protein to different time points across different observed nuclear volumes. From this, we found that both concentration of protein and K_OFF_ make a substantial contribution to the number of target site binding events (**Appendix SI, Fig. S8).** We then separated the contribution of changing amounts of CLOCK:BMAL1 protein and PER2:CRY1 mediated displacement to visits per minute by calculating the impact of removal of PER2.

We find that changing BMAL1 protein abundance accounts for the most variation in number of target site visitations, changing from 1.3 visits at the nadir to 2.9 visits at peak BMAL1 protein (**Fig. 4G**). CLOCK:BMAL1 mobility is supported by the action of PER2:CRY1 across all time points, accounting for a maximum of 26% visits at T32 and contributing to 10% of visits at the trough PER2 protein levels. To explore this relationship further, we tested the impact of altering the levels of PER2 in the stochastic model, choosing four PER2 concentrations, ranging from absent to greater than observed physiological levels (0, 10, 20 and 50 nM) (**Fig. 4H**). In the complete absence of PER2, BMAL1 mobility is hampered so that it visits less than 3 sites per minute. When PER2 spans the physiological range and beyond, a strong relationship in the visits per minute is forecast, rising by ~33%. Our modelling demonstrates dual modes of action of PER2:CRY1, repression via displacement of CLOCK:BMAL1 from target sites and facilitation of CLOCK:BMAL1 mobility to promote new target site binding (**Fig. 4I**). In this sense, PER2 acts both as part of a transcriptional repressor complex and as a facilitator of CLOCK:BMAL1 mobility to bind new target sites (34).

## Discussion

The circadian molecular circuit responds to and modulates an extraordinary number of biological processes, broadly imparted through DNA binding of CLOCK:BMAL1 to E-box sites (9). Through live cell microscopy of fluorescent ectopically and endogenously expressed circadian proteins we have sought to understand how the autonomous molecular clock regulates CLOCK:BMAL1 binding to DNA.

### Protein abundance and stoichiometry of the circadian circuit

Mathematical modelling demonstrates that low molecular abundances, as observed for core circadian components, lead to rapid internal and cell-to-cell desynchrony, which may be compensated for by strict control of stoichiometries and interactions (35). In the first instance, protein concentrations of both activators and repressors exert significant influence on amplitude as well as robustness of daily DNA binding cycles. We found approximately 16,000 BMAL1 and 8,000 CRY1 proteins per nucleus, consistent with our earlier reports for PER2 which found 12,000 proteins per nucleus in skin fibroblasts (11). In contrast with our doubling of endogenous CRY1 expression over PER2, a recent study by Gabriel et al. found approximately an eight-fold difference using the U20S, osteocarinoma cancer cell line, highlighting how different cell types and cell lines may diverge and influence the circadian network (14). Similarly, we observed significant disparities of endogenous BMAL1 across a range of cell types, with fibroblasts exhibiting a >2 fold increase in BMAL1 concentration when compared with chondrocytes (**SI appendix, Fig. S5**). The impact of cell type variation in protein concentrations and stoichiometries is difficult to discern but may confer tissue specific sensitives to clock control of output genes without the need for additional regulatory components, or could compensate the system, as evidenced by similar single site visitations despite a 4-fold decrease in nuclear volume (**Figure S8I**).

### Balance in affinity between BMAL1 and CLOCK may facilitate crosstalk

CLOCK was found to be cytoplasmic when ectopically expressed without BMAL1, with nuclear localization restored upon addition of BMAL1. This suggests BMAL1 oscillations could affect availability of nuclear CLOCK, consistent with several studies (17, 36). Here we found a 2:1 quantitative relationship suggesting a mechanism for maintenance for stoichiometry within the nucleus. It is worth noting that, despite a concentration of free BMAL1 of ~15 nM, only 10 percent of this was measured as part of the CLOCK:BMAL1 heterodimer. Low availability of heterodimeric transcription factor for DNA interactions, when compared with free protein, severely limits the potential to bind DNA, yet this is consistent with allowing other interactions to occur, including those reported with Hypoxia-inducible factor (HIF) and Aryl hydrocarbon receptor (AhR) (37, 38). Balancing availability of monomeric BMAL1 and CLOCK may therefore enable crosstalk with other pathways, or modulate interactions that have different affinities for monomeric versus heterodimeric CLOCK:BMAL1, as has been reported for CRY1 (8, 39).

### Impact of cycling CRY1 concentration, binding affinities and mode of repression on the clock

Substantive evidence for direct repression of BMAL1 transactivation by CRY1 now exists (8, 40). Here we have shown in live cells that this interaction is not only rhythmic but remarkably strong, with a higher affinity than any other protein pairings we have measured. This strong repression of CLOCK:BMAL1 by CRY1 balances against its low abundance. When acting without PER2, CRY1 exhibits persistent repression over 24 hours, likely owing to its regulated interaction with CLOCK and BMAL1, as evidenced by modelling the effect of removal of either cycling BMAL1, CRY1 or binding affinity between the two (**Fig. S4, SI appendix, Fig. S8**). This cycle in affinity provides evidence that the mammalian circadian clock also relies on oscillations in the ability of key proteins to heterodimerize one another. The exact mechanisms underlying this regulation of affinity are yet to be determined but could be hypothesized to be an outcome of dimerization with another partner that hinders or aids binding to CLOCK:BMAL1, such as PER2, or post-translational modifications leading to changes in affinity with BMAL1 (28, 30, 41). A ~25% shift in the diffusion of CRY1 equating to a change in mass close to that of, and in phase with the peak of, PER2 hints at the former proposition but further study is required (**Fig 2G** and **Fig 3E**).

### Individual genes exhibit a range of residence times

We found an average short residence time of 3 seconds for CLOCK:BMAL1, similar to other DNA binding transcription factors including GR, p53, p65 and STAT1 (19), potentially optimized to reduce gene expression noise (42). Here we modelled CLOCK:BMAL1 binding to a number of sites using an average off rate resulting in all sites behaving the same and demonstrating how DNA binding is globally regulated, in contrast with evidence from ChiP-seq, whereby different sites are differentially bound (9). Presumably, robustly detected peaks found by ChIP-seq represent genes with a slow unbinding rate, such as the E-box sites found in the DBP gene, which is supported by previous live cell imaging characterizing a longer 8 second residence time for BMAL1 on a DBP E-box concatemer (16). Altering the unbinding rates leads to a non-linear scaling in the occupation frequency (**SI appendix, Fig. S8**), highlighting the importance of regulating this parameter through post-translational modifications such as via phosphorylation of the CLOCK:BMAL1 complex as reported by Qin et al (43). Residence time may be tuned individually for different genes to ensure optimal transactivation, especially when considering recruitment of critical co-factors which do not interact with CLOCK:BMAL1 outside of DNA, as the probability of co-occupation increases with binding time. Ultimately, a considerable temporal gulf exists between the elaboration of a circadian rhythm (days) with the time-scale of DNA binding (seconds), altered by the accumulation of protein (hours).

Daily changes in BMAL1 protein are moderate, remaining as high as 10,000 molecules per nucleus even at the nadir of expression, resulting in many non-transcriptionally productive interactions of CLOCK:BMAL1 with DNA throughout the circadian cycle; these interactions however may be important, contributing to pioneer factor activity and allowing others genes to activate at a different phase to BMAL1 protein levels (44, 45).

### Compromise between E-box visitations and occupancy via PER:CRY mediated displacement

Whereas CRY1 inhibits BMAL1 transactivation via binding and blocking productive interactions with transcriptional coactivators, PER:CRY complexes permit an alternative mode of repression (8, 34). We demonstrate that increasing PER:CRY leads to an overall reduction in the ability for CLOCK: BMAL1 to remain bound through direct dimerization and manipulation of DNA unbinding. Work by Cao and Wang et al revealed PER2 removes CLOCK:BMAL1 in a CRY-dependent manner from E-Boxes via recruitment of CK1 and subsequent phosphorylation of CLOCK, effectively reducing affinity for DNA (34). Displacive repression of this kind reduces residency time on DNA sites and thus the number of sites bound at any one time. However, reducing residency time necessarily increases the rate at which a limited pool of transcription factors can move onto new sites, hence increasing the likelihood any one gene to be bound and reducing possible cell-to-cell variation. Site-specific residence times, most likely due to cofactor recruitment or chromatin modifications, coupled with this phenomenon would permit some gene targets to exhibit maximal CLOCK:BMAL1 binding beyond the time of the global peak. This supports findings by Menet and colleagues, who highlight groups of genes that have maximal binding events, as determined via ChIP-seq, outside of the zenith of total genome CLOCK:BMAL1 binding (44). We find that at the nadir of DNA binding, in the absence of PER:CRY, the visitations per minute would reduce by a quarter, whereas at the height of displacive repression CLOCK:BMAL1 is most mobile and thus more able to utilize site-specific binding factors. Furthermore, evidence of CLOCK:BMAL1 behaving as a so-called ‘kamikaze’ transcription factor, a factor most transcriptionally potent when phosphorylated and marked for degradation, implies that in addition to an increase in visitations per minute, transcriptional potency is also upregulated (16). Therefore, despite the relatively high efficiency of CLOCK:BMAL1 binding to DNA, it may spend much of its life performing transcriptionally non-productive tasks until modified via complexes such as PER:CRY.

## Materials and Methods

See supplementary material

## Acknowledgments

Supported by the Biotechnology and Biological Sciences Research Council, UK (awards BB/P017347/1 and BB/P017355/1 to ASIL and MHH), MRC core funding (MC_U105170643 to MHH), the National Institutes of Health (award GM107069 and GM141849 to CLP) and a University of California Office of the President Chancellor’s Postdoctoral Fellowship (to JLF). ASIL is a Wellcome Investigator (Grant 107851/Z/15/Z). AAK is supported by a Wellcome 4 year PhD studentship (216416/Z/19/Z).

## References

1. U. Bhadra, N. Thakkar, P. Das, M. Pal Bhadra, Evolution of circadian rhythms: from bacteria to human. Sleep Med 35, 49–61 (2017).

2. J. Gibbs et al., An epithelial circadian clock controls pulmonary inflammation and glucocorticoid action. Nat Med 20, 919–926 (2014).

3. N. Huang et al., Crystal structure of the heterodimeric CLOCK:BMAL1 transcriptional activator complex. Science 337, 189–194 (2012).

4. N. Gekakis et al., Role of the CLOCK protein in the mammalian circadian mechanism. Science 280, 1564–1569 (1998).

5. E. D. Buhr, J. S. Takahashi, Molecular components of the Mammalian circadian clock. Handb Exp Pharmacol 10.1007/978-3-642-25950-0_1, 3–27 (2013).

6. A. C. Liu et al., Redundant function of REV-ERBalpha and beta and non-essential role for Bmal1 cycling in transcriptional regulation of intracellular circadian rhythms. PLoS Genet 4, e1000023 (2008).

7. Y. Y. Chiou et al., Mammalian Period represses and de-represses transcription by displacing CLOCK-BMAL1 from promoters in a Cryptochrome-dependent manner. Proc Natl Acad Sci U S A 113, E6072–E6079 (2016).

8. H. Xu et al., Cryptochrome 1 regulates the circadian clock through dynamic interactions with the BMAL1 C terminus. Nat Struct Mol Biol 22, 476–484 (2015).

9. N. Koike et al., Transcriptional Architecture and Chromatin Landscape of the Core Circadian Clock in Mammals. Science 338, 349–354 (2012).

10. R. P. Aryal et al., Macromolecular Assemblies of the Mammalian Circadian Clock. Mol Cell 67, 770–782 e776 (2017).

11. N. J. Smyllie et al., Visualizing and Quantifying Intracellular Behavior and Abundance of the Core Circadian Clock Protein PERIOD2. Curr Biol 26, 1880–1886 (2016).

12. N. Yang et al., Quantitative live imaging of Venus::BMAL1 in a mouse model reveals complex dynamics of the master circadian clock regulator. PLoS Genet 16, e1008729 (2020).

13. O. T. Shafer, M. Rosbash, J. W. Truman, Sequential nuclear accumulation of the clock proteins period and timeless in the pacemaker neurons of Drosophila melanogaster. J Neurosci 22, 5946–5954 (2002).

14. C. H. Gabriel et al., Live-cell imaging of circadian clock protein dynamics in CRISPR-generated knock-in cells. Nat Commun 12, 3796 (2021).

15. J. Bagnall et al., Quantitative dynamic imaging of immune cell signalling using lentiviral gene transfer. Integr Biol (Camb) 7, 713–725 (2015).

16. M. Stratmann, D. M. Suter, N. Molina, F. Naef, U. Schibler, Circadian Dbp transcription relies on highly dynamic BMAL1-CLOCK interaction with E boxes and requires the proteasome. Mol Cell 48, 277–287 (2012).

17. R. V. Kondratov et al., BMAL1-dependent circadian oscillation of nuclear CLOCK: posttranslational events induced by dimerization of transcriptional activators of the mammalian clock system. Genes Dev 17, 1921–1932 (2003).

18. B. L. Sprague, J. G. McNally, FRAP analysis of binding: proper and fitting. Trends Cell Biol 15, 84–91 (2005).

19. J. Hettich, J. C. M. Gebhardt, Transcription factor target site search and gene regulation in a background of unspecific binding sites. J Theor Biol 454, 91–101 (2018).

20. L. Yu et al., A Comprehensive Review of Fluorescence Correlation Spectroscopy. Frontiers in physics. 9 (2021).

21. N. Dross et al., Mapping eGFP oligomer mobility in living cell nuclei. PLoS One 4, e5041 (2009).

22. K. Tsekouras, A. P. Siegel, R. N. Day, S. Presse, Inferring diffusion dynamics from FCS in heterogeneous nuclear environments. Biophys J 109, 7–17 (2015).

23. J. W. Krieger et al., Imaging fluorescence (cross-) correlation spectroscopy in live cells and organisms. Nat Protoc 10, 1948–1974 (2015).

24. B. T. Bajar et al., Improving brightness and photostability of green and red fluorescent proteins for live cell imaging and FRET reporting. Sci Rep 6, 20889 (2016).

25. H. Bennett, E. Aguilar-Martinez, A. D. Adamson, CRISPR-mediated knock-in in the mouse embryo using long single stranded DNA donors synthesised by biotinylated PCR. Methods 191, 3–14 (2021).

26. S. H. Yoo et al., PERIOD2::LUCIFERASE real-time reporting of circadian dynamics reveals persistent circadian oscillations in mouse peripheral tissues. Proc Natl Acad Sci U S A 101, 5339–5346 (2004).

27. J. M. Fustin, J. S. O’Neill, M. H. Hastings, D. G. Hazlerigg, H. Dardente, Cry1 circadian phase in vitro: wrapped up with an E-box. J Biol Rhythms 24, 16–24 (2009).

28. J. L. Fribourgh et al., Dynamics at the serine loop underlie differential affinity of cryptochromes for CLOCK:BMAL1 to control circadian timing. Elife 9 (2020).

29. R. Ye et al., Dual modes of CLOCK:BMAL1 inhibition mediated by Cryptochrome and Period proteins in the mammalian circadian clock. Genes Dev 28, 1989–1998 (2014).

30. I. Schmalen et al., Interaction of circadian clock proteins CRY1 and PER2 is modulated by zinc binding and disulfide bond formation. Cell 157, 1203–1215 (2014).

31. J. Bagnall et al., Quantitative analysis of competitive cytokine signaling predicts tissue thresholds for the propagation of macrophage activation. Sci Signal 11 (2018).

32. J. R. Beytebiere et al., Tissue-specific BMAL1 cistromes reveal that rhythmic transcription is associated with rhythmic enhancer-enhancer interactions. Genes Dev 33, 294–309 (2019).

33. D. T. Gillespie, A general method for numerically simulating the stochastic time evolution of coupled chemical reactions. Journal of Computational Physics 22, 403–434 (1976).

34. X. Cao, Y. Yang, C. P. Selby, Z. Liu, A. Sancar, Molecular mechanism of the repressive phase of the mammalian circadian clock. Proc Natl Acad Sci U S A 118 (2021).

35. D. Gonze, A. Goldbeter, Circadian rhythms and molecular noise. Chaos 16, 026110 (2006).

36. I. Kwon et al., BMAL1 shuttling controls transactivation and degradation of the CLOCK/BMAL1 heterodimer. Mol Cell Biol 26, 7318–7330 (2006).

37. J. Bagnall et al., Tight control of hypoxia-inducible factor-alpha transient dynamics is essential for cell survival in hypoxia. J Biol Chem 289, 5549–5564 (2014).

38. C. Jaeger, S. A. Tischkau, Role of Aryl Hydrocarbon Receptor in Circadian Clock Disruption and Metabolic Dysfunction. Environ Health Insights 10, 133–141 (2016).

39. A. K. Michael et al., Formation of a repressive complex in the mammalian circadian clock is mediated by the secondary pocket of CRY1. Proc Natl Acad Sci U S A 114, 1560–1565 (2017).

40. C. L. Gustafson et al., A Slow Conformational Switch in the BMAL1 Transactivation Domain Modulates Circadian Rhythms. Mol Cell 66, 447–457 e447 (2017).

41. M. Akashi et al., A positive role for PERIOD in mammalian circadian gene expression. Cell Rep 7, 1056–1064 (2014).

42. E. Azpeitia, A. Wagner, Short Residence Times of DNA-Bound Transcription Factors Can Reduce Gene Expression Noise and Increase the Transmission of Information in a Gene Regulation System. Front Mol Biosci 7, 67 (2020).

43. X. Qin, T. Mori, Y. Zhang, C. H. Johnson, PER2 Differentially Regulates Clock Phosphorylation versus Transcription by Reciprocal Switching of CK1epsilon Activity. J Biol Rhythms 30, 206–216 (2015).

44. J. S. Menet, S. Pescatore, M. Rosbash, CLOCK:BMAL1 is a pioneer-like transcription factor. Genes Dev 28, 8–13 (2014).

45. S. Klemz et al., Protein phosphatase 4 controls circadian clock dynamics by modulating CLOCK/BMAL1 activity. Genes Dev 10.1101/gad.348622.121 (2021).

