## Supplementary information for "Quantification of circadian interactions and protein abundance defines a mechanism for operational stability of the circadian clock"

### Supporting Information Text

### Contents

**1 Plasmids** 3

**2 Primary cell isolates and cell lines** 3

**3 Confocal microscopy** 3

**4 Real-time population bioluminescence recordings** 3

**5 Single-molecule Fluorescence in-situ hybridization** 3

**6 Fluorescence Recovery After Photo-bleaching** 3

**7 Fluorescence Correlation Spectroscopy** 4

**11 Tables and figures** 9

### 1. Plasmids

A set of lentivirus transfer plasmids encoding fluorescent fusions of circadian proteins were generated utilizing the gateway cloning system as previously described (1). In brief, an initial entry vector was cloned, containing murine coding sequences for: *Bmal1* (NM_007489.4), *Clock* (NM_007715.6), *Cry1* (NM_007771.3), and *Per2* (NM_011066.3). These vectors were then recombined with a target destination vector containing a fluorescent protein sequence to generate a terminal lentivirus vector, in which expression is regulated from the constitutive ubiquitin ligase C promoter. The NLS::EGFP, BMAL1 L95E and BMAL1 V435R encoding plasmids were all purchased from VectorBuilder.

### 2. Primary cell isolates and cell lines

Fibroblasts were isolated from lungs of adult mice. Lung tissue was dissected and homogenized before collagenase-1A (1.5 mg/ml, Cat no. C2674) treatment for 2 hours. The cell suspension was then filtered using 40 µm cell strainers before plating into DMEM (Cat no. D6429) supplemented with 10% fetal bovine serum (HyClone), penicillin-streptomycin (10 U/ml) and amphotericin B (2.5 µg/ml). Media was refreshed every 2-3 days for one week before sub-culturing or experimentation. Cells were sub-cultured for a maximum of 4 passages. SCN slice cultures were prepared as previously described (2) and imaged after 2-3 days after preparation for confocal imaging, or kept for 7 days in culture prior to bioluminescence recording.

NIH/3T3 (ATCC® CRL-1658™) cells were cultured in DMEM supplemented with 10% fetal bovine serum (HyClone). Cells were passaged every 2-3 days, maintaining cells till passage 30. Lentivirus transduced derivatives of these cells were made using low passage cultures (6-12). Production of 3rd generation lentivirus and subsequent transduction of NIH/3T3 cells was carried out as previously described (1). Singly transduced cells are referred to with a superscript LV1 prefix before the transgene. Sequential transductions were carried out a minimum of two weeks later and derived cells are then termed LV2. Circadian synchronization of cells was achieved by stimulation with 200 nM dexamethasone (Sigma D4902) for 1 hour before PBS washes and then switched to fresh culture media.

### 3. Confocal microscopy

For 2D culture imaging experiments, cells were plated into 35 mm glass bottomed imaging dishes (Greiner Bio-one) at least 6 hours prior to imaging. Measurements were performed using either a ZEISS LSM880 or ZEISS LSM780 microscope equipped with a stage mounted incubator to maintain cells at 37°C in humidified 5% $CO_2$; fluorescence image capture was performed using either ZEN 2.1 SP3 FP2 or ZEN 2010b SP1 software respectively. Fluorescence samples were excited using the most appropriate lasers; making use of an Argon-Ion laser to produce 488 nm or 514 nm excitation or diode laser to produce 561 nm excitation. The appropriate emitted fluorescence spectra were then collected using Quasar GasP array detectors. All images were made using a FLUAR 40x NA 1.3 oil immersion objective. Nuclear volume recordings were made by collecting a z-stack of images at nyquist rate using a 1 airy unit pinhole diameter and then analyzing images using Imaris (version 7.4). Time-lapse imaging of SCN: fluorescence timelapse recordings of CRY1::mRuby3 in SCN organotypic slices were acquired using Zeiss LSM780/880 inverted confocal system (Zeiss), and maintained at 37°C. Samples were placed in air-tight glass-bottom dishes (Mattek). Images were acquired using 10x objective, 30 seconds scan time per frame, 2 frames per hour, for 6 days for longer time lapse or 60-70 hours for shorter time lapse.

### 4. Real-time population bioluminescence recordings

Lung fibroblasts were plated into 35 mm plastic tissue culture dishes (Corning). Cell media was replaced with a HEPES buffered and phenol free DMEM. Additionally, D-luciferin was supplemented into the media 4-24 hours prior to recordings. To prevent gas-exchange, dishes were sealed with grease applied around the edges of the coverslips. Bioluminescence was then recorded by photomultiplier tubes (PMTs; Hamatasu) housed in an enclosed incubator at 37°C and without $CO_2$ as described previously (3).

### 5. Single-molecule Fluorescence in-situ hybridization

*Clock* and *Bmal1* mRNA were visualized using custom probes designed against *Clock* and *Bmal1* murine coding sequences via the Stellaris FISH Probe Designer (Biosearch Technologies Inc.). *Clock* and *Bmal1* probes were labeled with the Quasar-570 and Quasar-670 dyes respectively. Samples were imaged with a wide-field DeltaVision microscope as previously described and spot counting was performed with FISH-quant (4, 5).

### 6. Fluorescence Recovery After Photo-bleaching

FRAP was performed by time-lapse imaging of cells prior to and after photobleaching to visualize fluorescence recovery. Photobleaching of EGFP and Venus signals was performed using 488 nm or 514 nm laser lines respectively using circular regions of 5 µm diameter (approximately 10% nuclear area) and wholly within the nuclei of cells. Images were recorded every 0.262 seconds for up to 60 seconds.

We have used FRAP to infer DNA unbinding rates for BMAL1, see Fig. S2. The principle of the approach assumes that a combination of diffusion and binding to an unseen immobile substrate affects the speed in which fluorescent proteins move into the recovery area. Different trajectories of recovery therefore inform how the balance between binding and diffusion contributes to the apparent diffusion of the observed molecule. This approach has been utilized many times to characterize the binding times of transcription factors to DNA, including GR, STAT1, p53 and p65 and has been additionally cross validated against single molecule imaging (6). For our data, the recovery curves of BMAL1::EGFP (co-expressed alongside tagRFP::CLOCK) remained consistent when bleaching different sized nuclear regions indicating that binding contributes to the recovery profile rather than a solely diffusion-led process (Fig. S2B). For all subsequent measurements, a circular bleach region was used that was kept consistent across cells and accounted for approximately 10% of nuclear area. FRAP was performed and analyzed using the appropriate ZEN software version with recovery curves from the bleached region normalised to total cell fluorescence as well as background fluorescence (empty spaces away from cells). The normalised recovery curve of fluorescence within the bleached region over time, $t$, was then fit with a reaction binding model

$$f(t) = I_E - I_1 \mathrm{e}^{-\frac{t}{\tau}}, \quad [1]$$

where $\tau$ is the residence time (reciprocal of unbinding rate $k_{\mathrm{OFF}}$), $I_E$ and $I_1$ is the immobile and mobile fraction respectively (7).

### 7. Fluorescence Correlation Spectroscopy

**A. Experimental setup.** FCS measurements were performed in each cell nucleus using acquisition times of 20 seconds and a collection volume of 1 airy unit (approximately 0.722 fl and 1.10 for 488 nm and 561 nm excitation respectively for FLUAR 40x NA 1.3 oil immersions objective) calibrated in the x-y plane for maximum signal intensity. The effective confocal volumes were calculated via the equations

$$V_{\mathrm{eff}} = (2\pi)^{\frac{3}{2}} w_{xy}^2 w_z = (2\pi)^{\frac{3}{2}} \left(\frac{0.61\lambda}{\mathrm{NA}}\right)^2 \left(\frac{2n\lambda}{\mathrm{NA}^2}\right), \quad [2]$$

where $w_{xy}$ and $w_z$ is the beam width in the $x - y$ and $z$ directions respectively, with NA as the numerical aperture (NA = 1.3 for our 40x objective), $\lambda$ the wavelength of exciting laser and $n$ the refractive index of the immersion oil ($n = 1.515$ in all experiments). The appropriate spectra were collected for each different fluorophore. Laser power was reduced to minimize photo-bleaching whilst maintaining counts per molecule greater than 0.3 kHz.

**B. Fitting.** Auto-correlation curves extracted from the Zeiss .fcs files were fit over two rounds using a program written in Python 3; first a global parameter fit executed using a genetic algorithm *differential evolution* (SciPy (8)) generating initial guesses within reasonable parameter bounds was performed, followed by a final stage of non-linear least-squares regression implemented via the *curve fit* (SciPy) package with an arctan loss function. The non-linear regression was regularized using the standard deviation following the calculations by Saffarian and Elson (9) which incorporates systematic sources of error at short and long lag times due to the multi-tau correlation algorithm used to compute the correlation curve; at short lag times the averaging introduces uncertainty whilst at the long lag times less data points exist to correlate due to the finite time over which the experiment was run.

**C. Model selection.** The Akaike Information Criterion (AIC) (10) was used to score and select the best fit model with the lowest score, defined as

$$\mathrm{AIC} = 2k - 2\ln(\hat{L}), \quad [3]$$

where $k$ is the number of fitted parameters and $\hat{L}$ the maximum likelihood, equal to the sum of squared errors when using non-linear least squares regression to fit the curves. Results of the model selection for all FCS data sets performed in this study can be found in supplementary Figure S3.

**D. Interactions: Fluorescence Cross-Correlation Spectroscopy.** Care was taken for fluorescence cross-correlation spectroscopy (FCCS) measurements to avoid the green channel signal spilling up into the red channel causing false cross-correlation by reducing the laser power and observing the far-red part of the second channel. Control measurements were performed by selectively turning off either 488 or 561 nm lasers and tuning the red channel spectra until there was no cross-correlation due to spill-over. We analyzed both sets of auto-correlation and cross-correlation curves from the same measurement and used the same procedure as Sadaie et al 2014 (11) to calculate the disassociation constant $K_D$. We again used non-linear least squares regression upon this data, fitting the function

$$\frac{[\mathrm{Complex}]}{[\mathrm{A::GFP}]_{\mathrm{TOTAL}}} = \frac{[\mathrm{B::RFP}]_{\mathrm{TOTAL}} - [\mathrm{Complex}]}{K_D + [\mathrm{B::RFP}]_{\mathrm{TOTAL}} - [\mathrm{Complex}]}, \quad [4]$$

where $[\mathrm{A::GFP}]_{\mathrm{TOTAL}}$ is the total concentration of the protein $A$ fused to a green or yellow fluorescent protein, $[\mathrm{B::RFP}]_{\mathrm{TOTAL}}$ is the total of protein $B$ fused to a red fluorophore and [Complex] is the concentration of the dimer of $A$ and $B$ proteins. The standard deviation upon $K_D$ was also provided by this algorithm.

**E. Maturation correction.** Fluorescent proteins may take minutes or hours to fold correctly before becoming visible, with the invisible fraction becoming substantial if the degradation rate of the protein is comparable to the maturation rate, hence leading to misreports in protein number as measured by FCS. The red fluorescent protein, mRuby3 is known to have a long maturation time of 2.28 h (12) and CRY1 to have a half-life of approximately 2.1 h (13), therefore we applied a scaling correction to CRY1::mRuby3 FCS concentration data. To account for the unseen portion, we model the protein in two states; an invisible state, $P$, and a mature visible fraction, $M$. Assuming a constant rate of production, $k_p$, for the immature protein, a maturation rate for the fluorophore of $k_m$, and a degradation rate for both protein states of $k_d$ we get the set of ordinary differential equations

$$\frac{dP}{dt} = k_p - k_d P - k_m P, \qquad \frac{dM}{dt} = k_m P - k_d M. \qquad [5]$$

These equations may be solved analytically using an integrating factor assuming zero of both protein states at $t = 0$ and so long as the rate constants $k_m$ and $k_d$ are greater than zero. The unknown production rate, $k_p$, is divided out when computing the ratio of both states by $M$ and taking the limit of the solution as $t \longrightarrow \infty$ to yield the correction factor

$$c = \lim_{t\to\infty}\left(\frac{P(t)+M(t)}{M(t)}\right) = \frac{k_d + k_m}{k_m} = \frac{\tau_m}{\tau_d} + 1, \qquad [6]$$

where $\tau_m$ and $\tau_d$ are the doubling-time and half-life of the maturation and degradation respectively. Using equation (6), the half-life for CRY1 and the maturation time of mRuby3 we find a multiplicative factor of $c = 2.083$, which may multiply the observed protein to yield the total concentration of CRY1::mRuby3.

**F. Diffusion rate as a function of mass.** When considering normal diffusion due to Brownian motion the diffusion rate, $D$, may be modelled using the Stokes–Einstein equation (14)

$$D = \frac{k_B T}{8\pi\eta r}, \qquad [7]$$

where $k_B$ is the Boltzmann constant, $T$ the temperature in kelvin, $\eta$ the dynamic viscosity, and $r$ as the radius of the diffusing molecule. Assuming a constant density of spatially equally distributed constituent amino acids, the mass of the molecule grows like $r^3$ and hence the diffusion rate will be related to the mass of the molecule by

$$D \propto m^{-1/3}, \qquad [8]$$

hence a halving in mass will equate to an approximate increase of 1.26 times the diffusion rate.

### 8. In vitro binding assays

**A. Expression and purification of recombinant proteins.** Biotin Acceptor Peptide (BAP)-tagged CLOCK PAS-AB (mouse CLOCK residues 93-395) was expressed as a $His_6$-NusA-XL-tagged protein in *Escherichia coli* (*E. coli*) Rosetta2 (DE3) cells. The *E. coli* biotin ligase BirA was expressed as a GST-tagged protein in BL21 (DE3) cells. Protein expression was induced with 0.5 mM isopropyl-$\beta$-D-thiogalactopyranoside (IPTG) at an $OD_{600}$ of $\sim 0.8$ and grown for an additional 16 hours at 18°C. Cells were centrifuged at 4°C at 3200 x g, resuspended in 50 mM Tris pH 7.5, 300 mM NaCl, 5% (vol/vol) glycerol, and 5 mM $\beta$-mercaptoethanol (BME) and lysed using a microfluidizer followed by brief sonication on ice. After clarifying lysate by centrifugation at 4°C at 140,500 x g for 1 hour, proteins were captured using Ni-NTA resin (Qiagen) or Glutathione Sepharose 4B resin (GE Life Sciences). After extensive washing in in 50 mM Tris pH 7.5, 300 mM NaCl, 5% (vol/vol) glycerol, and 5 mM BME, the affinity and solubility tags (e.g. $His_6$-NusA-XL or GST) were cleaved on resin using GST-TEV or His6-TEV protease at 4°C overnight. Cleaved proteins were collected from the flow-through; GST-BirA was further purified using size exclusion chromatography (SEC) on a Superdex75 column (GE Healthcare) in 50 mM Tris, pH 8.0, 300 mM NaCl, 1 mM dithiothreitol (DTT), and 5% (vol/vol) glycerol, while CLOCK PAS-AB was further purified using SEC in 20 mM HEPES pH 7.5, 125 mM NaCl, 5% (vol/vol) glycerol, and 2 mM Tris(2-carboxyethyl)phosphine (TCEP).

BMAL1 PAS-AB (mouse BMAL1 residues 136-441) was expressed in Sf9 suspension insect cells (Expression systems) as a GST-tagged protein using the baculovirus expression system. Sf9 cells were infected with P3 virus at $1.2 \times 10^6$ cells per milliliter and grown for 72 hours at 27°C before harvesting. Cells were resuspended in resuspension buffer (50 mM HEPES pH 7.5, 300 mM NaCl, 5% (vol/vol) glycerol, and 5 mM $\beta$-mercaptoethanol (BME)). Cells were lysed using a microfluidizer followed by brief sonication on ice. After clarifying lysate by centrifugation at 140,500 x g for 1 hour at 4°C, the lysate was bound in batch-mode to Glutathione Sepharose 4B resin (GE Healthcare), washed in resuspension buffer and eluted with 50 mM HEPES pH 7.5, 150 mM NaCl, 5% (vol/vol) glycerol, 5 mM BME, and 25 mM reduced glutathione. The protein was desalted into 50 mM HEPES pH 7, 150 mM NaCl, 5% (vol/vol) glycerol, and 5 mM BME using a HiTrap Desalting column (GE Healthcare) and incubated with GST-TEV protease overnight at 4°C. The cleaved GST-tag and GST-tagged TEV protease were removed by Glutathione Sepharose 4B chromatography (GE Healthcare) and the BMAL1 PAS-AB was further purified by SEC on a Superdex75 column (GE Healthcare) in 20 mM HEPES pH 7.5, 125 mM NaCl, 5% (vol/vol) glycerol, and 2 mM TCEP. For long-term storage, small aliquots of proteins were frozen in liquid nitrogen and stored at -70°C.

Biotinylation of CLOCK PAS-AB. For the biotinylation reaction, 100 µM BAP-CLOCK PAS-AB was incubated in 20 mM HEPES pH 7.5, 125 mM NaCl, 5% (vol/vol) glycerol, and 2 mM TCEP with 2 mM ATP, 1 µM GST-BirA, and 150 µM biotin at 4°C overnight. GST-BirA was removed after the reaction using Glutathione Sepharose 4B resin (GE Healthcare) and excess biotin was separated from the labeled protein by SEC on a Superdex75 column in 20 mM HEPES pH 7.5, 125 mM NaCl, 5% (vol/vol) glycerol, and 2 mM Tris(2-carboxyethyl)phosphine (TCEP). Biotinylation of CLOCK PAS-AB was essentially complete, as determined by incubating the protein with excess streptavidin and resolving complexes on SDS-PAGE. For long-term storage, small aliquots of the biotinylated protein were frozen in liquid nitrogen and stored at -70°C.

**B. Surface plasmon resonance binding assays.** Kinetic binding experiments were conducted on a Biacore X100+ instrument (GE Healthcare) capturing biotinylated CLOCK PAS-AB on a streptavidin-coated SA sensor chip at 100-150 Response Units (RUs). Serial dilutions of BMAL1 PAS-AB from 0.25 to 10 nM were injected in phosphate buffered saline (PBS) over 250 seconds and dissociated into buffer over 250 seconds to determine binding kinetics. Sensorgram data were globally fit to a 1:1 biomolecular binding model with Biacore Evaluation software X100+ version 2.0.1 (GE Healthcare) to determine $k_{\mathrm{ON}}$, $k_{\mathrm{OFF}}$ and $K_D$. $\chi^2$ values $< 1$ and $R_{\max} \leq 100$ were established as quality cutoffs for acceptable data. See Figure S4F for surface plasmon resonance results between BMAL1 PAS-AB and CLOCK PAS-AB domains.

### 9. Animal lines

A previously established Venus::BMAL1 mouse line was used (15). Additionally, two more mouse lines were generated which included CRY1::mRuby3 (Fig S5) and a subsequent cross with mice expressing Venus::BMAL1 and PER2::LUC (16). The CRY1::mRuby3 mice were made using a CRISPR mediated genomic editing approach to introduce a fluorescent sequence via homology-directed repair. Details of methodology and validation of animals can be found in the supplementary materials. Eight to ten week old mice were housed in individual cages in light-tight cabinets (Tecniplast), equipped with activity mouse wheel cages (Actimetrics). Activity was recorded by ClockLab data collection software in 6-minute bins (Actimetrics). The mice were maintained at LD cycles (light on at 7 am; light off at 7 pm) for two weeks. Activity profiles were generated using ClockLab (Actimetrics) and used to apply Non-Parametric Circadian Rhythm Analysis (NPCRA) to 10 circadian days of wheel-running data, as described previously (17), to calculate: Intra-daily Variability (IV): Non-parametric frequency of activity-rest transitions within a day, with a range of between 0-2 (e.g. a Sine wave would have a value of 0 and Gaussian noise would have a value of 2). Inter-daily Stability (IS): Matching of activity patterns on day-to-day basis, ranging from 0 (Gaussian noise) to 1 (high-stability). Robust behavioral activity is characterized by low IV and high IS. ClockLab (Actimetrics) was used to generate double-plotted actograms with onsets of activity and phase angle of entrainment was calculated from 10 days of wheel-running data measuring the difference in time of the point in the entraining cycle (lights on) against the onset of activity.

**A. Generation of CRY1::mRuby3 mouse line.** We used CRISPR-Cas9 to generate C terminally tagged alleles for Cry1, see Fig. S6. Two sgRNA targeting the STOP codon of the gene were selected using the Sanger WTSI website (18) that adhered to our criteria for off target predictions (guides with mismatch (MM) of 0, 1 or 2 for elsewhere in the genome were discounted, and MM3 were tolerated if predicted off targets were not exonic). sgRNA sequences, wih PAM site indicated in italics, (aactgatacggtaaatactt-AGG and cggcagagcagtaactgata-CGG) were purchased as crRNA oligos, which were annealed with tracrRNA (both oligos supplied by IDT) in sterile, RNase free injection buffer (TrisHCl 1mM, pH 7.5, EDTA 0.1mM) by combining 2.5 mg crRNA with 5 mg tracrRNA and heating to 95°C, which was allowed to slowly cool to room temperature.

For our donor repair template we used the EASI-CRISPR long-ssDNA strategy (19), which comprised of the mRuby3 gene with linker flanked by 132 and 143 nt homology arms synthesized by a Biotinylation PCR and on-column denaturation method (20) (Fig. 2A). For embryo microinjection the annealed sgRNA was complexed with Cas9 protein (New England Biolabs) at room temperature for 10 minutes, before addition of long ssDNA donor (final concentrations; sgRNA 20 ng/ml, Cas9 protein 20 ng/ml, lssDNA 10 ng/ml). CRISPR reagents were directly microinjected into C57BL6/J (Envigo) zygote pronuclei using standard protocols. Zygotes were cultured overnight and the resulting 2 cell embryos surgically implanted into the oviduct of day 0.5 post-coitum pseudopregnant mice. Potential founder mice were screened by PCR (Fig. S6A), using primers that flank the sgRNA sites (Cut test F tacactatgctcacggggac and Cut test R accacgtcctcttcagaacc), which both identifies editing activity in the form of InDels from NHEJ repair, and can also detect larger products indicating HDR (Fig. S6A). Pups 18, 19 and 22, which gave positive products in PCR reactions, were sequenced by amplifying again with the cut test F/R primers using high fidelity Phusion polymerase (NEB), gel extracted and subcloned into pCRblunt (Invitrogen) and Sanger sequenced with M13 Forward and Reverse primers. All pups showed perfect sequence integration and were bred with a WT C57BL6/J to confirm germline transmission.

### 10. Mathematical modeling

**A. Modeling aims and assumptions.** We sought to model how the core circadian transcription factor, CLOCK:BMAL1, binds to specific E-BOX DNA sites over daily cycles in concentration and interactions with the key repressors CRYPTOCHROME1 (CRY1) and PERIOD2 (PER2). We have opted to neglect explicitly modeling the non-specific DNA interactions, such as sliding, hopping and intersegmental transfer, as we have no direct measurements of these properties. Instead we chose to allow the specific site on rate ($k_{\mathrm{ON}}$) to represent all protein-DNA processes required to achieve binding to an specific site by fitting $k_{\mathrm{ON}}$ alongside other ON rates. CRY1 and PER2 repress the activity of BMAL1 through direct binding of the transactivation domain (TAD) to block transcriptional potential and the promotion of weaker binding to DNA respectively. We have assumed that PER2 may interact with BMAL1 and CLOCK:BMAL1 only via CRY1 with the same affinity that CRY1 alone has for BMAL1. To constrain the on rates during fitting, except for CLOCK:BMAL1:CRY1:PER2 onto DNA represented by $R_{\mathrm{ON}}$, we have used the measured disassociation constants, $K_D$, between CLOCK-BMAL1 (Figure 1K), CLOCK:BMAL1-EBOX (21), CRY1-PER2 (Figure 3C) and the rhythmic CRY1-BMAL1 (Figure 2J). All protein-protein and protein-DNA interactions are modeled as explicit dimerization events leading to a new species dependent on an ON and OFF rate. A summary of the parameters, $K_D$ values and which parameters were proposed during during fitting is given in Table S2. Following the convention when defining chemical reactions, square brackets are used to signify concentrations of the species within. To aid understanding of the reactions being modeled we describe the species participating in reactions as familiar initializations, for example [CB] represents the concentration of the CLOCK:BMAL1 heterodimer and [C1] for CRY1. Consequently, further dimerizations or bound states are denoted by concatenations of these initializations, e.g. [CBC1P2] for the CLOCK:BMAL1:CRY1:PER2 tetramer or [CBS] for CLOCK:BMAL1 bound to a specific DNA ($S$) site.

**B. Dimerization.** Hetero-dimerization of two species $[A]$ and $[B]$ proceeds to the dimer $[AB]$ via the reaction

$$[A] + [B] \underset{k_{\mathrm{OFF}}}{\overset{k_{\mathrm{ON}}}{\rightleftharpoons}} [AB], \quad [9]$$

where where $k_{\mathrm{ON}}$ ($\mathrm{nM}^{-1}\,\mathrm{s}^{-1}$) and $k_{\mathrm{OFF}}$ ($\mathrm{s}^{-1}$) are the forward and backwards rates respectively (11), often referred to as the association and dissociation rate constants. In equilibrium the forward rate of reaction is equal to the backward rate resulting in the definition of the disassociation constant

$$K_D = \frac{[A][B]}{[AB]}, \quad [10]$$

defined in terms of the ON and OFF rates as

$$K_D = \frac{k_{\mathrm{OFF}}}{k_{\mathrm{ON}}}. \quad [11]$$

A stronger interaction is represented as a smaller $K_D$ value as the rate to the disassociated is smaller than the association rate. In the limit of long times, $t \to \infty$, the concentration of the dimer $[AB]$ in equilibrium becomes

$$[AB]_{\mathrm{eq}} = \frac{[A]_0 + [B]_0 + K_d - \sqrt{([A]_0 + [B]_0 + K_d)^2 - 4[A]_0[B]_0}}{2}, \quad [12]$$

where a subscript 0 denotes the initial concentration. Alternatively, assuming no production or degradation terms exist we may simulate analytically intractable multiple interactions by simulating a coupled ODE model until equilibrium concentrations are reached. For all ODE modeling we defined equilibrium as less than a 1% deviation in molecular concentrations over the last 20% of simulated time points. In all cases equilibrium was established in less than 30 minutes of simulated time, smaller than the window over which experimental FCS time point measurements were performed.

**C. Ordinary differential equation model of DNA binding.** Systems of ordinary differential equations (ODEs), modeling the concentrations of molecular species, were solved in the Python 3 programming language to reflect measured interactions between different molecules and DNA. All interactions are modeled as explicit dimerization events yielding a new molecular species. ODEs were solved in time as an initial value problem using the LSODA solver as implemented in the SciPy *odeint* function (8) and ran until equilibrium concentrations were reached, typically reached in less than 30 minutes of simulated time. The system of ODEs are

$$
\begin{aligned}
\frac{d[CB]}{dt} &= -d_{\mathrm{ON}}[CB][C1P2] + d_{\mathrm{OFF}}[CBC1P2] - d_{\mathrm{ON}}[CB][C1] + d_{\mathrm{OFF}}[CBC1] - k_{\mathrm{ON}}[CB][S] + k_{\mathrm{OFF}}[CBS] \\
&\quad + b_{\mathrm{ON}}[C][B] - b_{\mathrm{OFF}}[CB], \\
\frac{d[S]}{dt} &= -k_{\mathrm{ON}}[CB][S] + k_{\mathrm{OFF}}[CBS] - R_{\mathrm{ON}}[CBC1P2][S] + R_{\mathrm{OFF}}[CBC1P2S] - k_{\mathrm{ON}}[CBC1][S] + k_{\mathrm{OFF}}[CBC1S], \\
\frac{d[CB]}{dt} &= -d_{\mathrm{ON}}[CB][C1P2] + d_{\mathrm{OFF}}[CBC1P2] - d_{\mathrm{ON}}[CB][C1] + d_{\mathrm{OFF}}[CBC1] - k_{\mathrm{ON}}[CB][S] + k_{\mathrm{OFF}}[CBS] \\
&\quad + b_{\mathrm{ON}}[C][B] - b_{\mathrm{OFF}}[CB], \\
\frac{d[S]}{dt} &= -k_{\mathrm{ON}}[CB][S] + k_{\mathrm{OFF}}[CBS] - R_{\mathrm{ON}}[CBC1P2][S] + R_{\mathrm{OFF}}[CBC1P2S] - k_{\mathrm{ON}}[CBC1][S] + k_{\mathrm{OFF}}[CBC1S], \\
\frac{d[CBS]}{dt} &= k_{\mathrm{ON}}[CB][S] - k_{\mathrm{OFF}}[CBS] - d_{\mathrm{ON}}[CBS][C1] + d_{\mathrm{OFF}}[CBC1S] - d_{\mathrm{ON}}[CBS][C1P2] + d_{\mathrm{OFF}}[CBC1P2S], \\
\frac{d[C1]}{dt} &= -d_{\mathrm{ON}}[CB][C1] + d_{\mathrm{OFF}}[CBC1] - d_{\mathrm{ON}}[CBS][C1] + d_{\mathrm{OFF}}[CBC1S] - a_{\mathrm{ON}}[C1][P2] + a_{\mathrm{OFF}}[C1P2] \\
&\quad - d_{\mathrm{ON}}[B][C1] + d_{\mathrm{OFF}}[BC1], \\
\frac{d[CBC1]}{dt} &= -a_{\mathrm{ON}}[CBC1][P2] + a_{\mathrm{OFF}}[CBC1P2] + d_{\mathrm{ON}}[CB][C1] - d_{\mathrm{OFF}}[CBC1] - k_{\mathrm{ON}}[CBC1][S] + k_{\mathrm{OFF}}[CBC1S] \\
&\quad + b_{\mathrm{ON}}[C][BC1] - b_{\mathrm{OFF}}[CBC1], \\
\frac{d[CBC1S]}{dt} &= d_{\mathrm{ON}}[CBS][C1] - d_{\mathrm{OFF}}[CBC1S] - a_{\mathrm{ON}}[CBC1S][P2] + a_{\mathrm{OFF}}[CBC1P2S] + k_{\mathrm{ON}}[CBC1][S] - k_{\mathrm{OFF}}[CBC1S], \\
\frac{d[P2]}{dt} &= -a_{\mathrm{ON}}[CBC1][P2] + a_{\mathrm{OFF}}[CBC1P2] - a_{\mathrm{ON}}[C1][P2] + a_{\mathrm{OFF}}[C1P2] - a_{\mathrm{ON}}[CBC1S][P2] + a_{\mathrm{OFF}}[CBC1P2S] \\
&\quad - a_{\mathrm{ON}}[BC1][P2] + a_{\mathrm{OFF}}[BC1P2], \\
\frac{d[C1P2]}{dt} &= -d_{\mathrm{ON}}[CB][C1P2] + d_{\mathrm{OFF}}[CBC1P2] + a_{\mathrm{ON}}[C1][P2] - a_{\mathrm{OFF}}[C1P2] - d_{\mathrm{ON}}[CBS][C1P2] + d_{\mathrm{OFF}}[CBC1P2S] \\
&\quad - d_{\mathrm{ON}}[B][C1P2] + d_{\mathrm{OFF}}[BC1P2], \\
\frac{d[CBC1P2S]}{dt} &= R_{\mathrm{ON}}[CBC1P2][S] - R_{\mathrm{OFF}}[CBC1P2S] + a_{\mathrm{ON}}[CBC1S][P2] - a_{\mathrm{OFF}}[CBC1P2S] \\
&\quad + d_{\mathrm{ON}}[CBS][C1P2] - d_{\mathrm{OFF}}[CBC1P2S], \\
\frac{d[CBC1P2]}{dt} &= a_{\mathrm{ON}}[CBC1][P2] - a_{\mathrm{OFF}}[CBC1P2] + d_{\mathrm{ON}}[CB][C1P2] - d_{\mathrm{OFF}}[CBC1P2] - R_{\mathrm{ON}}[CBC1P2][S] \\
&\quad + R_{\mathrm{OFF}}[CBC1P2S] + b_{\mathrm{ON}}[C][BC1P2] - b_{\mathrm{OFF}}[CBC1P2], \\
\frac{d[C]}{dt} &= -b_{\mathrm{ON}}[C][B] + b_{\mathrm{OFF}}[CB] - b_{\mathrm{ON}}[C][BC1] + b_{\mathrm{OFF}}[CBC1] - b_{\mathrm{ON}}[C][BC1P2] + b_{\mathrm{OFF}}[CBC1P2], \\
\frac{d[B]}{dt} &= -b_{\mathrm{ON}}[C][B] + b_{\mathrm{OFF}}[CB] - d_{\mathrm{ON}}[B][C1] + d_{\mathrm{OFF}}[BC1] - d_{\mathrm{ON}}[B][C1P2] + d_{\mathrm{OFF}}[BC1P2], \\
\frac{d[BC1]}{dt} &= d_{\mathrm{ON}}[B][C1] - d_{\mathrm{OFF}}[BC1] - b_{\mathrm{ON}}[C][BC1] + b_{\mathrm{OFF}}[CBC1] - a_{\mathrm{ON}}BC1[P2] + a_{\mathrm{OFF}}[BC1P2], \\
\frac{d[BC1P2]}{dt} &= d_{\mathrm{ON}}[B][C1P2] - d_{\mathrm{OFF}}[BC1P2] - b_{\mathrm{ON}}[C][BC1P2] + b_{\mathrm{OFF}}[CBC1P2] + a_{\mathrm{ON}}[BC1][P2] - a_{\mathrm{OFF}}[BC1P2].
\end{aligned}
$$

A genetic algorithm, implemented in *differential evolution* SciPy (8)), was utilized to fit the unknown parameters in the ODE model via Chi-squared minimization to experimental $k_{\mathrm{OFF}}$ mean and standard error on the mean using an *In silico* value, $\bar{k}_{OFF}$, generated by the model. All non-dimerized species concentrations, as measured experimentally, were introduced for each of the seven time-points – a 24-hour time span sampled every 4 hours – into the model as inputs alongside measured disassociation constants to constrain fitted OFF rates as a function of proposed ON rates, reducing the number of fitted parameters. A summary of the parameters in the model is given in Table S2. During fitting the *In silico* $k_{\mathrm{OFF}}$ value was calculated by allowing all species to reach equilibrium after setting all DNA bound species to zero following an initial run of the model, the resultant equilibrium concentrations of bound and free molecules were used to calculate the off rate (Figure S8A). The average apparent DNA unbinding rate $\bar{k}_{OFF}$, which is analogous to the same rate as experimentally measured in FRAP, is simulated following the method by Röding et al (22) through rearranging equation (11) for the off rate

$$
\bar{k}_{OFF} = k_{\mathrm{ON}} K_D = k_{\mathrm{ON}} \frac{[\text{Unbound CB}][\text{Unbound Sites}]}{[\text{Bound Sites}]}, \qquad [13]
$$

with

$$[\text{Unbound CB}] = [CB]_{\text{eq}} + [CBC1]_{\text{eq}} + [CBC1P2]_{\text{eq}}, \quad [14]$$

$$[\text{Unbound Sites}] = [S]_{\text{eq}}, \quad [15]$$

$$[\text{Bound sites}] = [CBS]_{\text{eq}} + [CBC1S]_{\text{eq}} + [CBC1P2S]_{\text{eq}} \quad [16]$$

where eq denotes concentrations at equilibrium as $t \to \infty$. The apparent $\bar{k}_{OFF}$ is an average of the CLOCK:BMAL1-DNA binding OFF rate $k_{\text{OFF}}$ and CLOCK:BMAL1:CRY1:PER2-DNA OFF rate $R_{\text{OFF}}$ weighted by their respective relative concentrations, with increasing levels of CRY1:PER2 increasing $\bar{k}_{OFF}$ as $k_{\text{OFF}} < R_{\text{OFF}}$. The fitted parameters are given in in Table S2 and predicted *In silico* $\bar{k}_{OFF}$ values for WT and PER2 KO can be seen in Figure S8C. Knocking out PER2 (keeping all other species and parameters the same as wild type values) removes all rhythmic regulation of $\bar{k}_{OFF}$ and ensures that CLOCK:BMAL1 is bound for longer at all time points such that the number of bound specific sites (S) also increases for all time points (Figure S8E). Locking BMAL1, CRY1, PER2 and the interaction between BMAL1 & CRY1 to their mean value between 24-48 hours post-dexamethosone (DEX) treatment, reveals that setting BMAL1 to its mean value significantly alters both bound and free from DNA CLOCK:BMAL1:CRY1 whilst locking the other rhythmic components has little impact, Figure S8F.

Upper and lower bounds on one-at-a-time (OAT) sensitivity analysis (Figure 4A-B) were generated by running the fitted model for both an estimate of the smallest and largest number of target sites, 1000 and 10,000 respectively, with the mean representing the mean number of target sites from ChIP data, namely 3436. We may estimate the number of target sites for CLOCK:BMAL1 from previous studies investigating high confidence sites that BMAL1 binds to in ChIP-seq, with table S1 outlining the reports and peaks measured via ChIP-seq that were used in estimating the number of target sites used in our mathematical modeling. In addition to the OAT analyses in Figure 4A-B we also examined how changing amounts of CRY1:PER2 alters the residence time of CLOCK:BMAL1 on DNA, demonstrating how CRY1:PER2 readily promotes removal from DNA in a non-linear fashion over a physiologically plausible range of concentrations (Figure S8B).

**D. Stochastic DNA binding model.** Stochastic binding simulations in Python 3 utilized the Gillespie algorithm (23) through the StochPy library (24) to simulate a reduced topology, considering only CLOCK:BMAL1 and CLOCK:BMAL1:CRY1 binding to sites with the addition of 1 extra marked site, $M$, and using the fitted ON/OFF rates from the ODE model. The table S3 gives the reactions and propensities that was modeled. Thirty runs over 60 minutes were used to generate mean and standard deviations with times to reach 95% of all sites at least once determined via fitting of an inverse exponential to the number of unique site visits counted via the variable $A_{CB}$. The time for available CLOCK:BMAL1 complexes to bind 95% of all binding sites at least once is calculated by fitting the recovery curve $f(t) = 1 - \exp(-\lambda t)$ to normalised stochastic trajectories of $S_{tot} - S$ ($S_{tot} = 3436$), see Figure S8H, and then converting the recovery rate $\lambda$ using the equation

$$\tau_{95\%} = \frac{\ln(20)}{\lambda}. \quad [17]$$

Visits per minute to a single site were calculated by counting binding and unbinding to $M$, which possesses the same ON and OFF rates as other target-sites. Assessment of the contribution of PER2 mediated displacement was performed by setting PER2 concentration to zero (KO) in the ODE model and using the simulated OFF rate in a parallel run to wild-type (WT) runs (Figure S8C), with the reduced number of visits attributed to the slower OFF rate. Furthermore, we observed the same behavior in this reduced stochastic model, when compared to the full ODE model, for PER2 KO as the mean and standard deviation of the number of sites bound by CLOCK:BMAL1 in both WT and KO conditions, Figure S8G, being comparable to the ODE model results in Figure S8E. Finally, to assess the differences that would be induced by different nuclear volumes, as seen between different cell types, we ran the stochastic model at the same molecular concentrations over two volumes; a small volume of 240 fl representative of a typical mouse embryonic fibroblast (MEF) or various immune cell types (see Figure S5E) and 926 fl as measured for our lung fibroblasts used throughout this study, Figure S8I. We note little difference in the rate at which CLOCK:BMAL1 visits the single marked site $M$, indicating that the increase in DNA sites comparatively to the number of molecules at a smaller nuclear volume was balanced by the increase in ON rate due to the now higher concentration of DNA.

### 11. Tables and figures

**Table S1. BMAL1 ChIP reports**

| No. | Tissue | BMAL1 peaks | Reference |
|---|---|---|---|
| 1 | Liver | 2049 | Rey et al 2011 (25) |
| 2 | Liver | 5952 | Koike et al 2012 (26) |
| 3 | U2OS | 2001 | Wu et al 2017 (27) |
| 4 | PECS | 2026 | Oishi et al 2017 (28) |
| 5 | Liver | 4813 | Beytebiere et al 2019 (29) |
| 6 | Kidney | 4034 | Beytebiere et al 2019 (29) |
| 7 | Heart | 2520 | Beytebiere et al 2019 (29) |
| 8 | NIH3T3 | 4740 | Chiou et al 2016 (30) |
| 9 | Skeletal muscle | 2787 | Dyar et al (31) |
| | Mean average | 3436 | |

**Table S2. Summary of ordinary differential equation model parameters**

* Fixed during fitting
† Proposal parameter
Model fit $\chi^2 = 7.53$

| No. | Parameter | Unit | Description | Fitted value (SD) | Disassociation constant ($\mathrm{nM}$) |
|---|---|---|---|---|---|
| 1 | $k_{\mathsf{ON}}$† | $\mathrm{nM}^{-1}\,\mathrm{s}^{-1}$ | CLOCK:BMAL1 DNA binding on rate | $(0.268 \pm 9.398) \times 10^{-1}$ | } $K_D = 10^{*}$ (21) |
| 2 | $k_{\mathsf{OFF}}$ | $\mathrm{s}^{-1}$ | Specific DNA site off rate | $0.268 \pm 9.398$ | |
| 3 | $b_{\mathsf{ON}}$† | $\mathrm{nM}^{-1}\,\mathrm{s}^{-1}$ | CLOCK - BMAL1 binding rate | $0.736 \pm 1.634$ | } $K_D = 147.6 \pm 9.8^{*}$ |
| 4 | $b_{\mathsf{OFF}}$ | $\mathrm{s}^{-1}$ | CLOCK - BMAL1 unbinding rate | $(1.086 \pm 2.413) \times 10^{2}$ | |
| 5 | $d_{\mathsf{ON}}$† | $\mathrm{nM}^{-1}\,\mathrm{s}^{-1}$ | BMAL1 - CRY1 binding rate | $1.160 \pm 3.496$ | } Variable $K_D^{*}$ see Fig. 2J |
| 6 | $d_{\mathsf{OFF}}$ | $\mathrm{nM}^{-1}\,\mathrm{s}^{-1}$ | BMAL1 - CRY1 unbinding rate | $d_{\mathsf{ON}} \times K_D$(BMAL1:CRY1) (Fig. 2J) | |
| 7 | $a_{\mathsf{ON}}$† | $\mathrm{nM}^{-1}\,\mathrm{s}^{-1}$ | PER2 - CRY1 binding rate | $7.798 \pm 2.718$ | } $K_D = 82 \pm 4.9^{*}$ |
| 8 | $a_{\mathsf{OFF}}$ | $\mathrm{s}^{-1}$ | PER2 - CRY1 unbinding rate | $(6.39 \pm 2.26) \times 10^{2}$ | |
| 9 | $R_{\mathsf{ON}}$† | $\mathrm{nM}^{-1}\,\mathrm{s}^{-1}$ | CLOCK:BMAL1:CRY1:PER2 DNA binding on rate | $(0.071 \pm 2.547)$ | } $K_D = 176.0$ |
| 10 | $R_{\mathsf{OFF}}$† | $\mathrm{s}^{-1}$ | CLOCK:BMAL1:CRY1:PER2 DNA off rate | $(1.253 \pm 0.591) \times 10^{1}$ | |

**Table S3. Stochastic model reactions and propensities**

Counter for arrivals by CLOCK:BMAL1 ($CB$) without CRY1 ($C1$) to previously unbound sites $S$ converting them to $S_0$ given by $A_{CB}$ as well as counters for marked site $M$ binding represented by $B_X$, and unbinding, $U_X$, by species $X$.

| No. | Reaction | Propensity |
|---|---|---|
| 1 | $CB + S \longrightarrow CBS + A_{CB}$ | $k_{\text{ON}} \cdot CB \cdot S$ |
| 2 | $CB + S_0 \longrightarrow CBS$ | $k_{\text{ON}} \cdot CB \cdot S_0$ |
| 3 | $CBS \longrightarrow CB + S_0$ | $k_{\text{OFF}} \cdot CBS$ |
| 4 | $CBC1 + S \longrightarrow CBC1S$ | $k_{\text{ON}} \cdot CBC1 \cdot S$ |
| 5 | $CBC1S \longrightarrow CBC1 + S$ | $k_{\text{OFF}} \cdot CBC1S$ |
| 6 | $CBC1 + S_0 \longrightarrow CBC1S_0$ | $k_{\text{ON}} \cdot CBC1 \cdot S_0$ |
| 7 | $CBC1S_0 \longrightarrow CBC1 + S_0$ | $k_{\text{OFF}} \cdot CBC1S_0$ |
| 8 | $CB + M \longrightarrow CBM + B_{CB}$ | $k_{\text{ON}} \cdot CB \cdot M$ |
| 9 | $CBM \longrightarrow CB + M + U_{CB}$ | $k_{\text{OFF}} \cdot CBM$ |
| 10 | $CBC1 + M \longrightarrow CBC1M + B_{CBC1}$ | $k_{\text{ON}} \cdot CBC1 \cdot M$ |
| 11 | $CBC1M \longrightarrow CBC1 + M + U_{CBC1}$ | $k_{\text{OFF}} \cdot CBC1M$ |

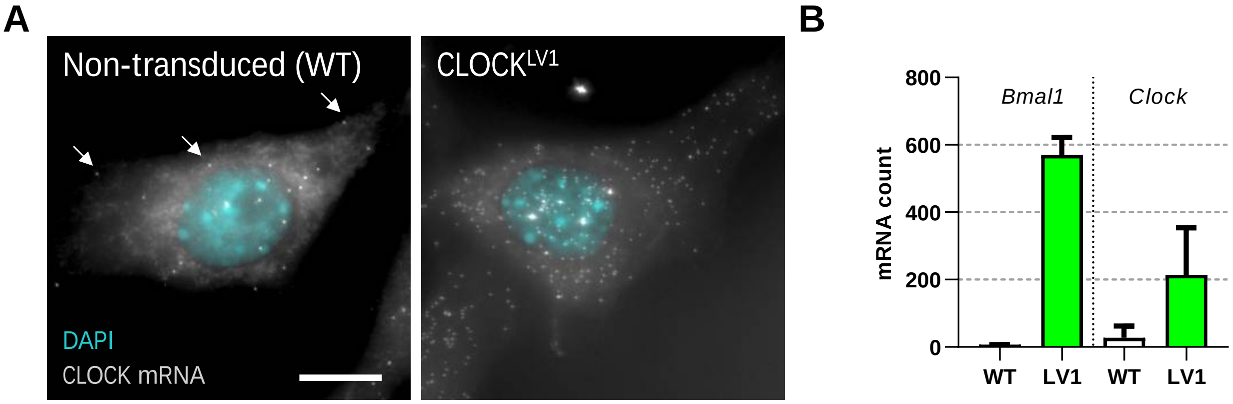

**Fig. S1. Ectopically expressed mRNA is the major form in a lentivirus transduced system** (A) NIH/3T3 cells were stained for CLOCK mRNA by single-molecule fluorescent in situ hybridisation (smFISH). (B) Mature mRNA was counted from images of many single cells to determine the mRNA content per cell for both non-transduced and cells transduced to express EGFP::CLOCK from a ubiquitin ligase C promoter (n=2177 cells for WT and n=155 cells for LV1). Increased nuclear bright dots can be seen for the transduced cells. These bright dots correspond to sites of transcription which is increased beyond two copies due to the multiple sites of integration following transduction.

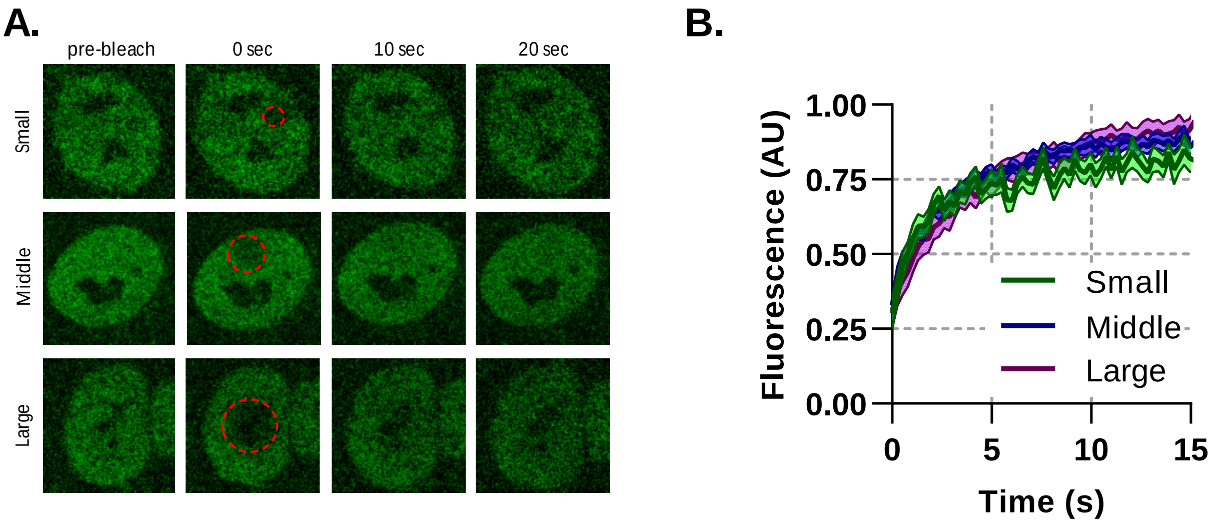

**Fig. S2. Binding plays a significant role in BMAL1 mobility** Fluorescence recovery after photobleaching for NIH/3T3 cells transduced with tagRFP::CLOCK and BMAL1:EGFP. (A) Imaging protocol was performed on BMAL1::EGFP signal. Regions of photobleaching are shown as a red-dotted line which is increased in size. (B) The bleached region recovery curves are shown as averages of all cells with an SEM error envelope (n=14, 20 and 15).

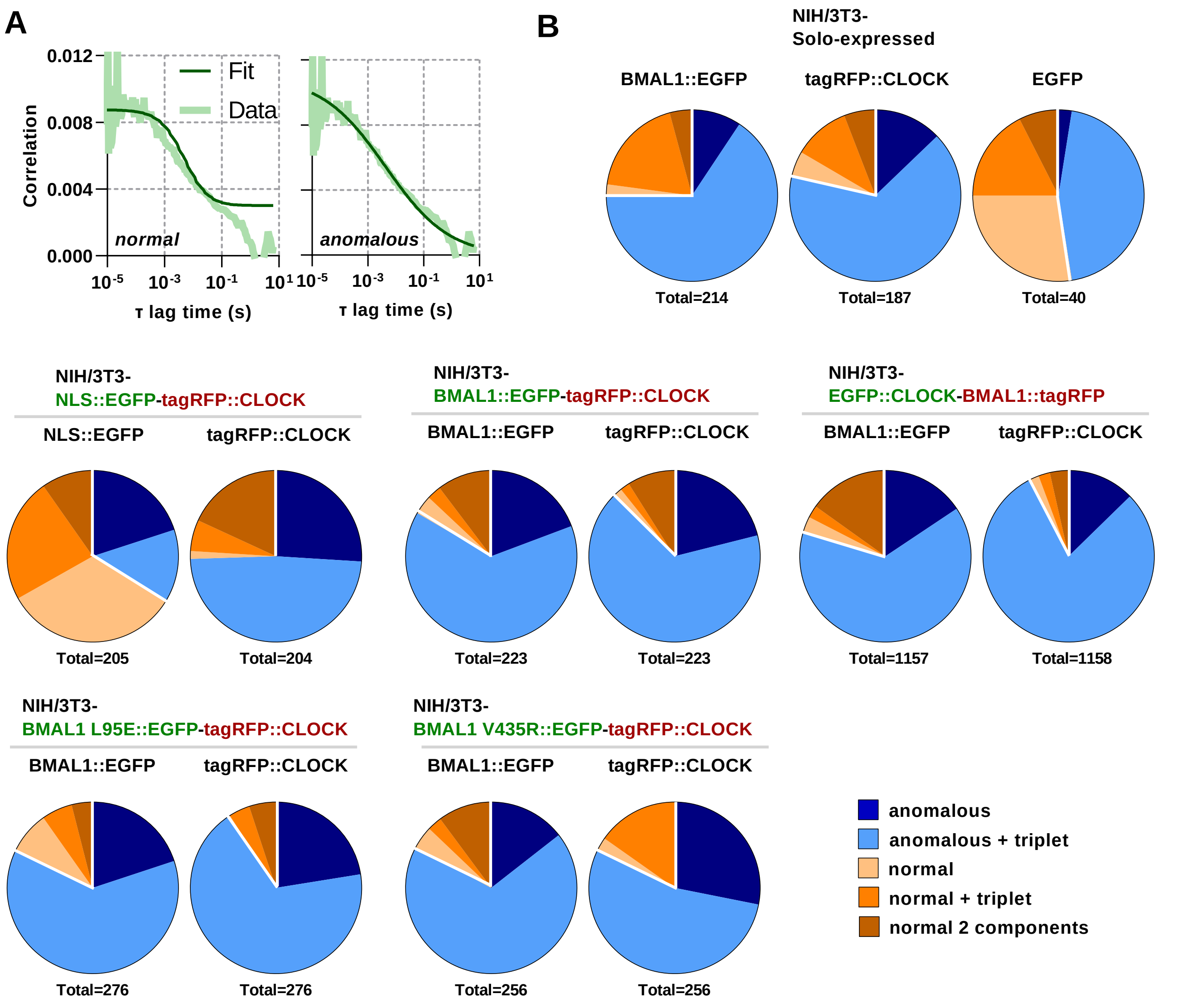

**Fig. S3. Anomalous diffusion best fits protein movement** (A) Representative normal and anomalous model fits for BMAL1::EGFP FCS data sets for cells transduced with lentivirus to express BMAL1, CLOCK or control fluorescent proteins. (B) Summary data sets for all five models considered. Each model was fit to each measurement and the fit with the lowest AIC score selected. A single pie chart was generated for cells carrying a single fluorescent label whereas two pie charts are plotted for cells with multiple labels, corresponding to the analyzed green or red fluorescence.

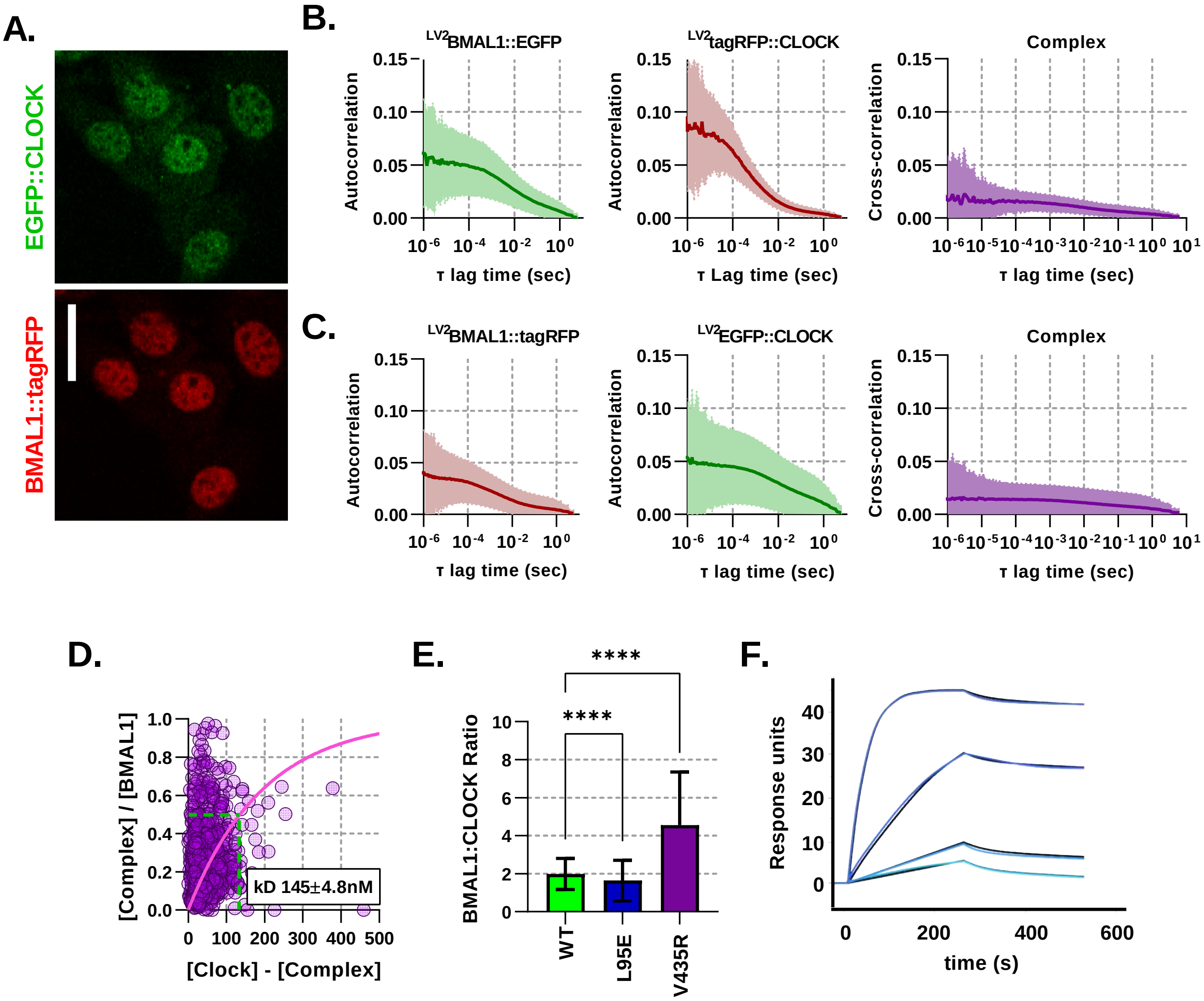

**Fig. S4. Fluorescent BMAL1 and CLOCK proteins behave similarly when colors are swapped** (A) Confocal images of NIH/3T3-LV2 EGFP::CLOCK-BMAL1::tagRFP cells. (B) Average auto- and cross- correlation curves shown as mean (line) and standard deviation (error envelope) for NIH/3T3 cells transduced to express BMAL1::EGFP and tagRFP::CLOCK. (C) Same as previous for color swapped cells so that they express BMAL1::tagRFP and EGFP::CLOCK (n=1158). (D) Dissociation plot to determine $K_D$ for data from C. (E) Ratio calculations for number of nuclear molecules of BMAL1::EGFP/NLS::EGFP to tagRFP::CLOCK. (F) Surface plasmon resonance (SPR) analysis of heterodimer formation with immobilized biotinylated CLOCK PAS-AB in the presence of increasing concentrations of BMAL1 PAS-AB from 0.25 to 10 nM (light to dark blue). Data were fitted using a 1:1 binding model (global fit in black, association rate $k_{\text{ON}} = 6.47 \times 10^5$ M-1s-1, dissociation rate $k_{\text{OFF}} = 8.98 \times 10^{-4}$ s-1, $K_D = 1.39 \times 10^{-9}$ M, $\chi^2 = 0.230$). Kruskal-Wallis test used to determine significance (values are denoted as p>0.05 ns, p<0.05 *, p<0.01 **, p<0.001 *** and p<0.0001 ****)

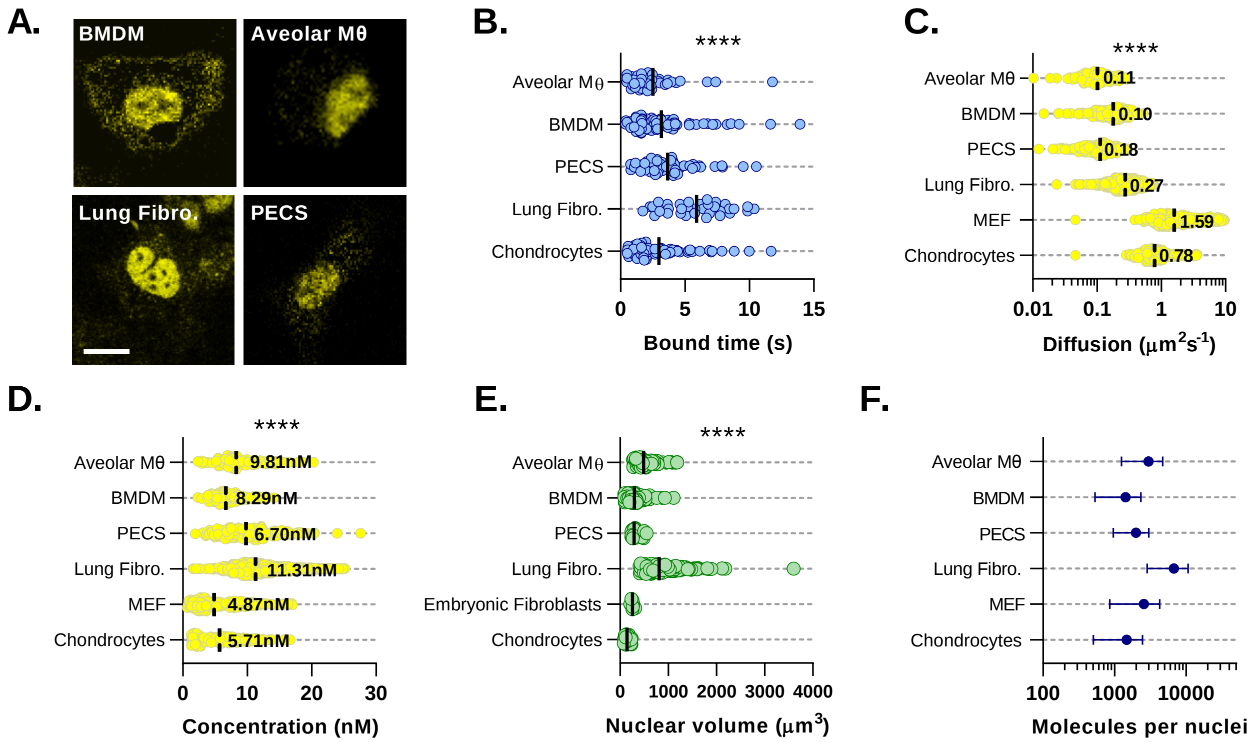

**Fig. S5. BMAL1 concentration and DNA binding parameters minimally vary across cell types** (A) Confocal microscopy images of primary cultures isolated from Venus::BMAL1 mice. (B) Characteristic bound time calculated from FRAP measurements of primary cultures from A. Chondrocyte data was previously measured by Yang et al. 2020 and reanalyzed for bound time (n = 73, 87, 61, 42 and 35 cells). (C) Cell cultures were measured using FCS and the auto-correlation data used to determine Venus::BMAL1 diffusion coefficient and (D) protein concentration (n = 107, 142, 156, 1597, 243 and 172 cells). (E) To determine total molecular abundance per nuclei, cultures were stained with Hoechst 33342 and then imaged. Nuclear volumes were then determined (n= 84, 115, 27, 169, 9 and 30). (F) Total BMAL1 molecules calculated from average nuclear concentration and average nuclear volume. Kruskal-Wallis test was used to determine significance (values are denoted as p>0.05 *, p<0.01 **, p<0.001 *** and p<0.0001 ****)

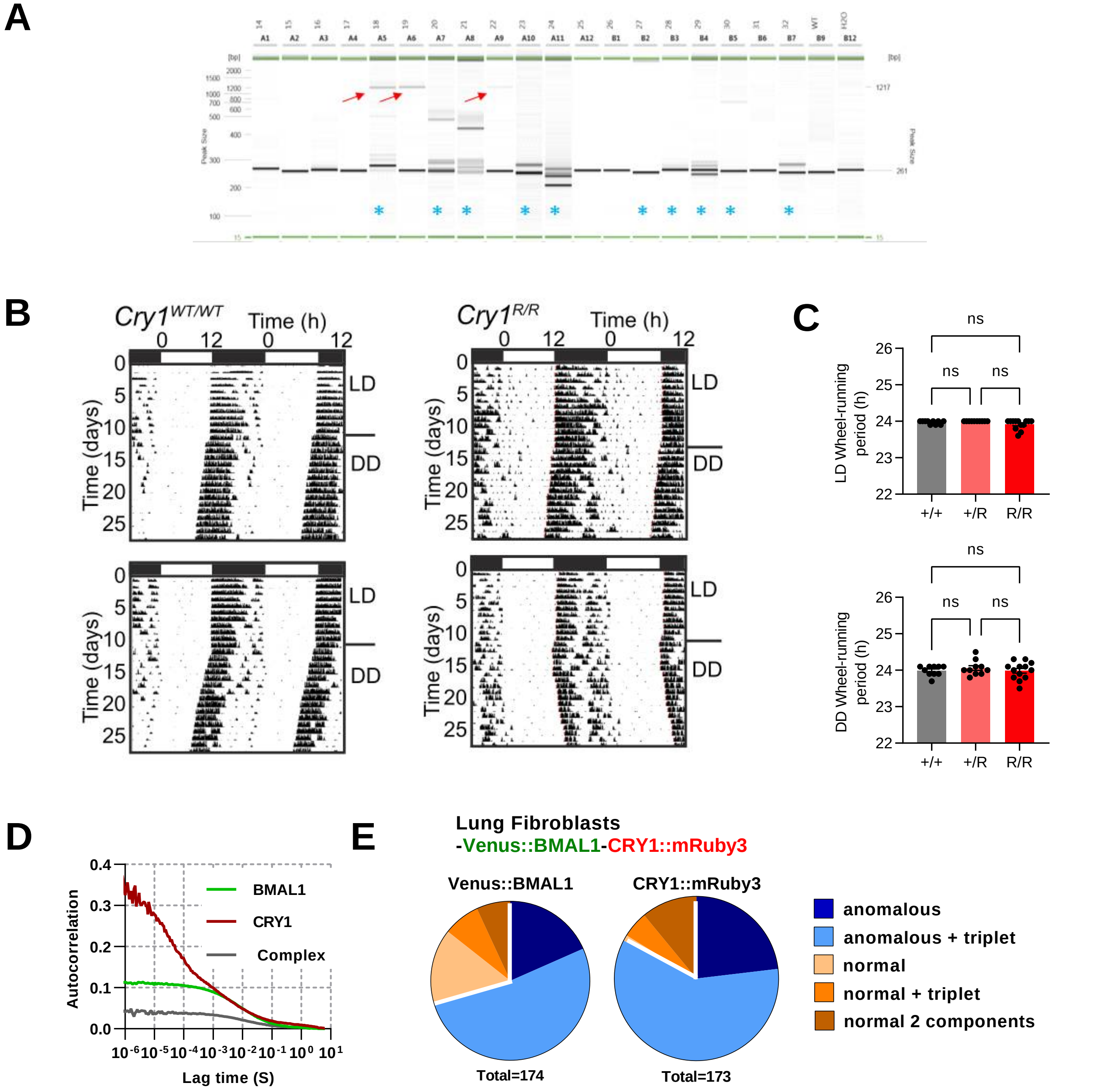

**Fig. S6. Generation of CRY1::mRuby3 mouse line** (A) Genotyping of pups by PCR. Images are PCR reactions run on Qiaxcel with red arrows indicating correct HDR product size, blue asterisks indicate mice in which InDels resulting from NHEJ are observed. (B) Actogram traces for wild-type and genetically modified CRY1::mRuby3 mice. (C) Mean ± SEM circadian periods for wheel-running in light-dark conditions (12h/12h) (+/+ = 10; +/R = 10; R/R = 13). Mean ± SEM circadian periods for wheel-running in constant dark (+/+ = 6; +/R = 10; R/R = 10). (D) Mean correlation curve for FCS measurements of BMAL1 x CRY1 x PER2::luc lung fibroblasts 24h after dexamethasone synchronization (n=144, BMAL1 and n=135, CRY1). (E) FCS model selection results (pooling 24 to 48 hours post-dexamethasone measurements). One-way ANOVA test used to determine significance (values are denoted as p>0.05 ns, p<0.05 *, p<0.01 **, p<0.001 *** and p<0.0001 ****)

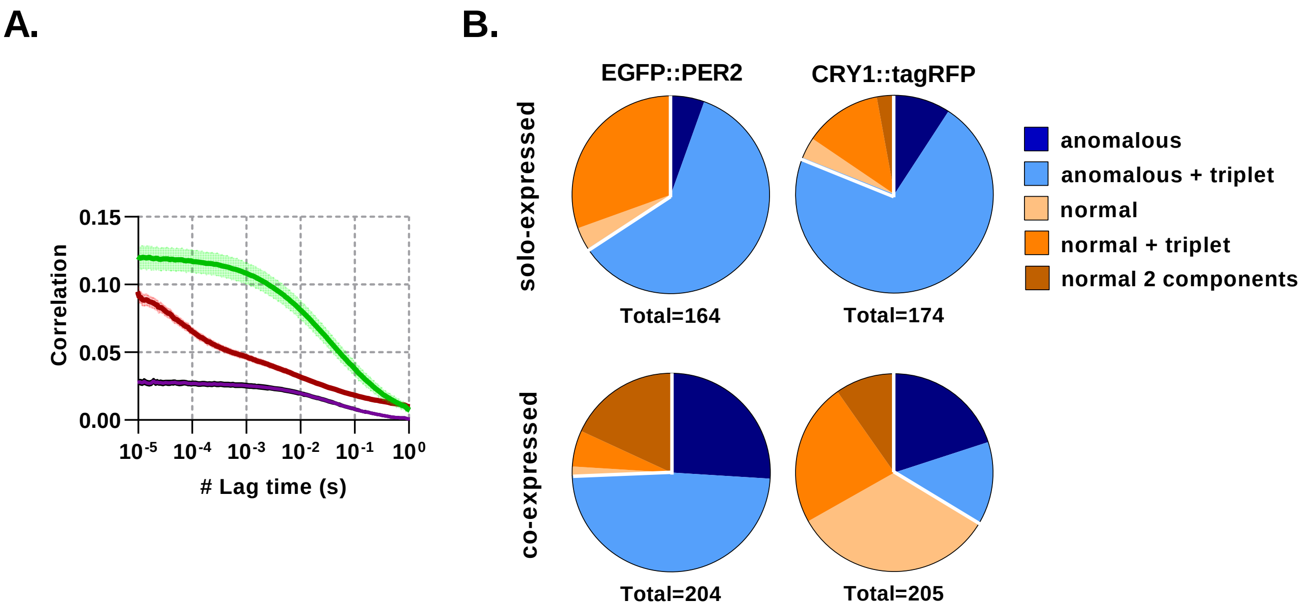

**Fig. S7. CRY1 mobility is affected by co-expression with PER2** (A) Average auto- and cross- correlation curves shown as mean (line) and standard deviation (error envelope) for NIH/3T3 cells transduced to express EGFP::PER2 and CRY1::tagRFP. Measurements were made in the nuclei. (B) FCS model selection results for cells that either solo-express CRY1/PER2 or co-express both.

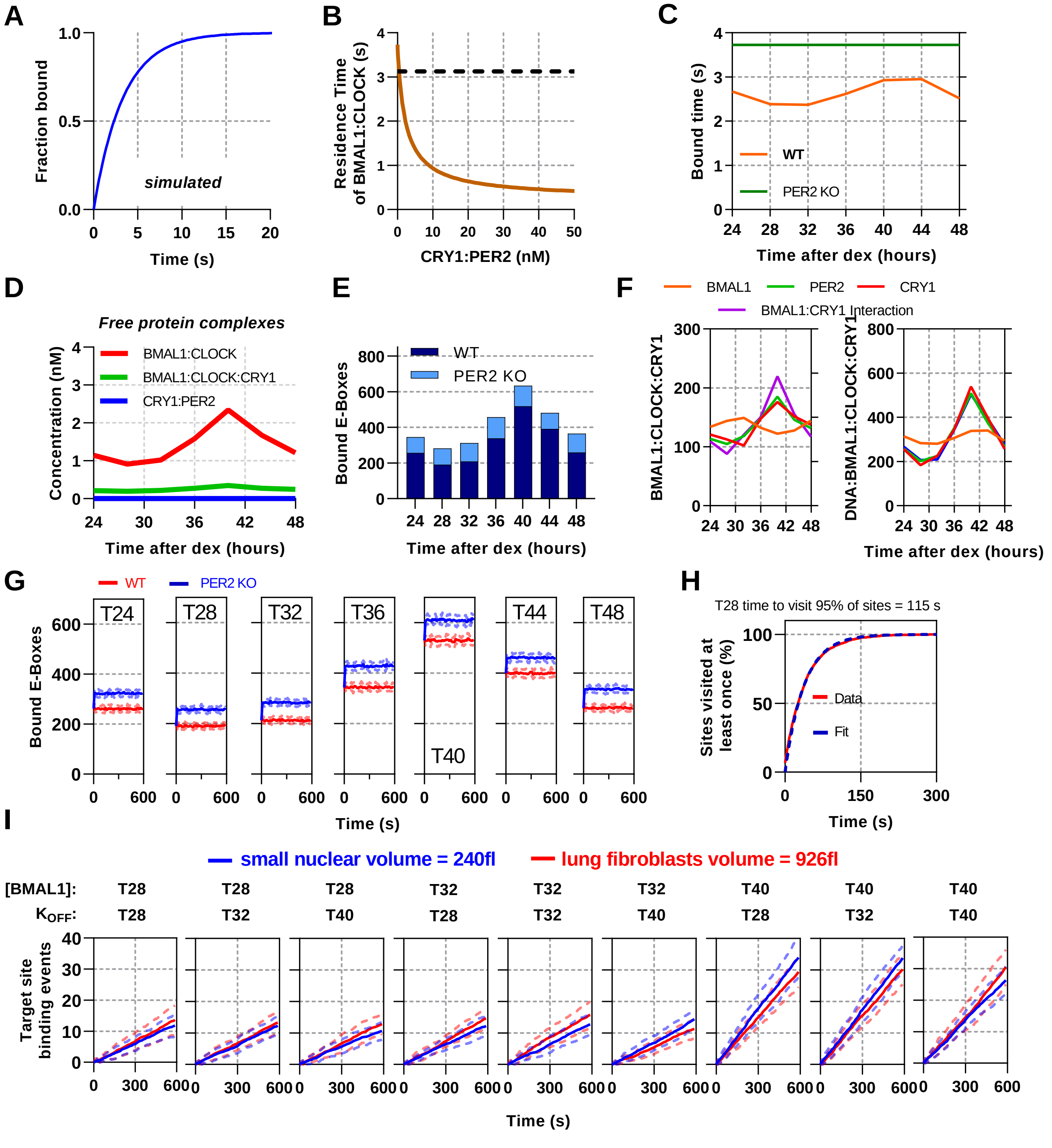

**Fig. S8. CLOCK:BMAL1 ODE and stochastic binding models over experimentally measured parameters** (A) Normalized model simulated FRAP using ODE model by counting recovery of site bound BMAL1 species after removal. (B) Model determined residence time of DNA bound CLOCK:BMAL1 across different concentrations of CRY1:PER2 using parameters from T40. (C) Modeled residence time of DNA bound CLOCK:BMAL1 across circadian time calculated for WT and without PER2. Without PER2 model data over a circadian cycle showing (D) non DNA-bound complexes and (E) DNA bound CLOCK:BMAL1. (F) Plots showing the CLOCK:BMAL1:CRY1 complex across several simulated conditions, including removal of rhythmicity from BMAL1, CRY1 or PER2 protein levels or the interaction between BMAL1:CRY1 (fixing them to their mean concentration). (G) Stochastic model showing the average binding (with SD) of CLOCK:BMAL1 bound target sites using input measurements from all time points for both WT and without PER2 simulations. (H) The time to visit every E-Box site once for T28 showing fit. (I) Model simulation plots showing CLOCK:BMAL1 visits to a single promoter over time. Red and blue lines show simulations using different nuclear volumes (red and blue lines, corresponding to primary lung fibroblasts and mouse embryonic fibroblasts). Input values for BMAL1 concentration and binding OFF rate correspond to those measured at the time point referred above each panel.

### References

1. J Bagnall, et al., Quantitative dynamic imaging of immune cell signalling using lentiviral gene transfer. *Integr. Biol.* **7**, 713–725 (2015).
2. N Smyllie, et al., Visualizing and quantifying intracellular behavior and abundance of the core circadian clock protein period2. *Curr. Biol.* **26**, 1880–1886 (2016).
3. A Loudon, et al., The biology of the circadian ck1 tau mutation in mice and syrian hamsters: A tale of two species. *Cold Spring Harb. Symp. on Quant. Biol.* **72**, 261–271 (2007).
4. J Bagnall, et al., Quantitative analysis of competitive cytokine signaling predicts tissue thresholds for the propagation of macrophage activation. *Sci. Signal.* **11** (2018).
5. F Mueller, et al., Fish-quant: automatic counting of transcripts in 3d fish images. *Nat. Methods* **10**, 277–278 (2013).
6. FL Groeneweg, et al., Quantitation of glucocorticoid receptor DNA-binding dynamics by single-molecule microscopy and FRAP. *PLoS ONE* **9** (2014).
7. BL Sprague, JG McNally, Frap analysis of binding: proper and fitting. *Trends Cell Biol.* **15**, 84–91 (2005).
8. P Virtanen, et al., SciPy 1.0: Fundamental Algorithms for Scientific Computing in Python. *Nat. Methods* **17**, 261–272 (2020).
9. S Saffarian, EL Elson, Statistical analysis of fluorescence correlation spectroscopy: The standard deviation and bias. *Biophys. J.* **84**, 2030–2042 (2003).
10. H Akaike, A new look at the statistical model identification. *IEEE Transactions on Autom. Control.* **19**, 716–723 (1974).
11. W Sadaie, Y Harada, M Matsuda, K Aoki, Quantitative in vivo fluorescence cross-correlation analyses highlight the importance of competitive effects in the regulation of protein-protein interactions. *Mol. Cell. Biol.* **34**, 3272–3290 (2014).
12. E Balleza, JM Kim, P Cluzel, Systematic characterization of maturation time of fluorescent proteins in living cells. *Nat. Methods* **15**, 47–51 (2017).
13. SH Yoo, et al., Competing e3 ubiquitin ligases govern circadian periodicity bydegradation of cry in nucleus and cytoplasm. *Cell* **152**, 1091—1105 (2013).
14. A Einstein, Über die von der molekularkinetischen theorie der wärme geforderte bewegung von in ruhenden flüssigkeiten suspendierten teilchen. *Annalen der Physik* **322**, 549–560 (1905).
15. N Yang, et al., Quantitative live imaging of venus::bmal1 in a mouse model reveals complex dynamics of the master circadian clock regulator. *PLOS Genet.* **16**, 1–24 (2020).
16. SH Yoo, et al., Period2::luciferase real-time reporting of circadian dynamics reveals persistent circadian oscillations in mouse peripheral tissues. *Proc. Natl. Acad. Sci.* **101**, 5339–5346 (2004).
17. SM Reppert, DR Weaver, Coordination of circadian timing in mammals. *Nature* **418**, 935–941 (2002).
18. A Hodgkins, et al., WGE: a CRISPR database for genome engineering. *Bioinformatics* **31**, 3078–3080 (2015).
19. RM Quadros, et al., Easi-crispr: a robust method for one-step generation of mice carrying conditional and insertion alleles using long ssdna donors and crispr ribonucleoproteins. *Genome Biol.* **18**, 92 (2017).
20. H Bennett, E Aguilar-Martinez, AD Adamson, Crispr-mediated knock-in in the mouse embryo using long single stranded dna donors synthesised by biotinylated pcr. *Methods* **191**, 3–14 (2021) Methods of genome engineering and model validation.
21. N Huang, et al., Crystal structure of the heterodimeric clock:bmal1 transcriptional activator complex. *Science* **337**, 189–194 (2012).
22. M Röding, L Lacroix, A Krona, T Gebäck, N Lorén, A highly accurate pixel-based frap model based on spectral-domain numerical methods. *Biophys. J.* **116**, 1348–1361 (2019).
23. DT Gillespie, Exact stochastic simulation of coupled chemical reactions. *The J. Phys. Chem.* **81**, 2340–2361 (1977).
24. TR Maarleveld, BG Olivier, FJ Bruggeman, Stochpy: A comprehensive, user-friendly tool for simulating stochastic biological processes. *PLOS ONE* **8**, 1–10 (2013).
25. G Rey, et al., Genome-wide and phase-specific dna-binding rhythms of bmal1 control circadian output functions in mouse liver. *PLOS Biol.* **9**, 1–18 (2011).
26. N Koike, et al., Transcriptional architecture and chromatin landscape of the core circadian clock in mammals. *Science* **338**, 349–354 (2012).
27. Y Wu, et al., Reciprocal regulation between the circadian clock and hypoxia signaling at the genome level in mammals. *Cell Metab.* **25**, 73–85 (2017).
28. Y Oishi, et al., Bmal1 regulates inflammatory responses in macrophages by modulating enhancer rna transcription. *Sci. Reports* **7**, 7086 (2017).
29. JR Beytebiere, et al., Tissue-specific bmal1 cistromes reveal that rhythmic transcription is associated with rhythmic enhancer–enhancer interactions. *Genes &; Dev.* **33**, 294–309 (2019).
30. YY Chiou, et al., Mammalian period represses and de-represses transcription by displacing clock–bmal1 from promoters in a cryptochrome-dependent manner. *Proc. Natl. Acad. Sci.* **113** (2016).
31. KA Dyar, et al., Transcriptional programming of lipid and amino acid metabolism by the skeletal muscle circadian clock. *PLOS Biol.* **16**, 1–47 (2018).
